# Foxi2 and Sox3 are master regulators controlling ectoderm germ layer specification

**DOI:** 10.1101/2025.01.09.632114

**Authors:** Clark L. Hendrickson, Ira L. Blitz, Amina Hussein, Kitt D. Paraiso, Jin Cho, Michael W. Klymkowsky, Matthew J. Kofron, Ken W.Y. Cho

## Abstract

In vertebrates, germ layer specification represents a critical transition where pluripotent cells acquire lineage-specific identities. We identify the maternal transcription factors Foxi2 and Sox3 to be pivotal master regulators of ectodermal germ layer specification in *Xenopus*. Ectopic co-expression of Foxi2 and Sox3 in prospective endodermal tissue induces the expression of ectodermal markers while suppressing mesendodermal markers. Transcriptomics analyses reveal that Foxi2 and Sox3 jointly and independently regulate hundreds of ectodermal target genes. During early cleavage stages, Foxi2 and Sox3 pre-bind to key cis-regulatory modules (CRMs), marking sites that later recruit Ep300 and facilitate H3K27ac deposition, thereby shaping the epigenetic landscape of the ectodermal genome. These CRMs are highly enriched within ectoderm-specific super-enhancers (SEs). Our findings highlight the pivotal role of ectodermal SE-associated CRMs in precise and robust ectodermal gene activation, establishing Foxi2 and Sox3 as central architects of ectodermal lineage specification.

**Highlights:** Foxi2 and Sox3 are master regulators for the ectodermal germ layer and sub-lineages

Five ectodermal cell states of early gastrulae are regulated by Foxi2 and Sox3

Foxi2 and Sox3 binding sites in the genome are enriched in ectoderm super-enhancers

SE-associated genes show high expression levels, low noise, stabilizing ectodermal regulation

## INTRODUCTION

A major breakthrough in biology was the first animal (*Xenopus laevis*) cloning experiment, which proved that differentiated cells retain the full genetic potential to develop into an entire organism (Gurdon et al., 1958). This result demonstrated the concept of genomic equivalence, where cellular differentiation is governed by gene regulation rather than by irreversible modifications to the genome, such as differential loss of subsets of genes in cell lineages. These experiments further revealed that the egg cytoplasm contains necessary maternal regulators to reprogram the epigenetic memory of the differentiated cell nuclei to a totipotent state. Subsequently it was shown in mammalian cells that networks of master regulator transcription factors (TFs), including most notably Klf4, Myc, Pou5f1, and Sox2, and are critical for embryonic pluripotency (Takahashi and Yamanaka, 2006). This highlights the essential role of multiple TFs working in concert to orchestrate cellular identity, lineage specification, and reprogramming.

Additionally, this discovery of these pluripotency TFs among various TF permutations highlights that not all TFs hold equally importance in development. Some TFs play a more critical role in regulating specific differentiation pathways because they occupy a higher position within a gene regulatory program controlling downstream events in differentiation and maintaining pluripotency. One of the earliest TFs to support the concept of master regulators of differentiation was Myod1, which was shown to regulate muscle cell differentiation (Davis et al., 1987). Other examples include Spi1 (PU.1), which specifies myeloid and B-cell lineages in hematopoiesis (Fisher and Scott 1998), Gata1 in erythroid lineage specification (Moriguchi and Yamamoto, 2007), and Pdx1 for pancreas organogenesis (Horb et al., 2003). These master regulatory TFs play a pivotal role in determining the identity and function of specific cell types. Identifying master regulatory TFs in developmental biology is fundamentally important for comprehending the mechanisms of cell lineage segregation, and advancing regenerative medicine to facilitate direct cell lineage conversion.

Our goal is to determine the primary master regulators initiating ectodermal cell lineage differentiation at the highest positions in the programming of cell lineges. In vertebrates, germ layer specification is one of the earliest developmental decisions that pluripotent cells make. In amphibians, the three germ layers are organized along the animal-vegetal axis, where the ectoderm forms in the animal pole, the endoderm vegetally, and the mesoderm in the equator. The ectoderm comprises progenitor cell populations giving rise to the skin, nervous system, sensory organs and the numerous neural crest derivatives. Several experiments have demonstrated that *Xenopus* ectodermal cells of the blastula stage are pluripotent. Transplantation of blastula prospective ectodermal cells into the vegetal endoderm alters the ectodermal cells to adopt endodermal fates (Snape et al., 1987). In addition, explanted blastula ectodermal tissue (“animal caps”) can be converted to mesendodermal cells *ex vivo* (organoids), by soaking them in activin, a TGFβ-superfamily ligand (Smith et al., 1990; Asashima et al.,1990; van den Eijnden-Van Raaij, et al., 1990). With specific combinations of growth factors and/or small molecules, blastula ectodermal explants can be programmed to differentiate into a wide range of cell types, including neural tissue, muscle cells, blood, heart and endoderm primordia (Ariizumi et al., 2017; Green et al., 1990). Although this ectodermal pluripotency persists through blastula stages, it is autonomously lost by early gastrula (Gurdon et al., 1985; Jones et al., 1987). Therefore, ectodermal germ layer and sub-lineage specification is an ideal system to study the dynamic processes controlling cellular potency and differentiation.

Earlier epigenetic studies have shown that the early embryonic genome lacks significant histone modifications before the blastula stage and therefore remains epigenetically naïve (reviewed in Blitz and Cho, 2021). Histone modifications first appear during blastula stage, coinciding with major zygotic genome activation (ZGA) and the segregation of the pluripotent embryonic cells into specialized ectoderm, mesoderm, and endodermal lineages. We hypothesize that localized expression of master TFs enables them to bind to specific cis-regulatory modules (CRMs), thereby activating or repressing the transcription of target genes, thus orchestrating the gene regulatory programs required for cell lineage specification.

Here we reveal that the ectodermal gene regulatory program is orchestrated by maternally expressed, and ectodermally enriched master regulatory TFs, Foxi2 and Sox3. These TFs prebind to the CRMs of the naïve embryonic genome prior to ZGA, preceding the appearance of major histone modifications and transcription. This prebinding event coordinates subsequent ectodermal cell differentiation program. The CRMs bound by Foxi2 and Sox3 are associated with ectodermal super enhancers (SEs), which ensure the robust target gene expression. Our findings are consistent with the view that Foxi2 and Sox3 as master regulators of the ectodermal germ layer.

## RESULTS

### Different epigenetic states accrue on ectoderm- and endoderm-expressed genes during ZGA

Epigenetic analyses of *Xenopus* blastula stage embryos show that the embryonic genome is largely devoid of major histone tail modifications such as H3K27ac, H3K27me3, and H3K4me1 (Gupta et al., 2014, van Heeringen et al., 2014). These marks begin to accrue during blastula stages, becoming widespread in the genome during gastrulation (van Heeringen et al., 2014; Hontelez et al., 2015; Charney et al., 2017; Paraiso et al., 2019). However, a limitation of whole-embryo epigenetic experiments, which include cells from all three germ layers, is the inability to determine whether these histone modifications occur in a spatially restricted manner.

To gain spatial information about histone modification states in different germ layers, early gastrulae (stage 10.5) were dissected into ectoderm and endoderm fragments approximately 3 hours after the onset of ZGA (stage 8.5) (Fig S1A, B). Histone modifications H3K27ac and H3K27me3 were then examined (Fig. S1A, B) around enhancers and promoters to understand their role in regulating the transcription of associated genes. Using early embryonic time course (Owens et al. 2016), and early gastrula embryo dissection (Blitz et al. 2017) RNA-seq data, we determined the top 250 zygotically expressed genes specifically enriched in either the ectoderm or endoderm (Table S1). The germ layer specific histone mark deposition of H3K27ac and H3K27me3 was plotted within a 25kb region upstream and downstream of the genes (Figure 1A). The mesoderm was excluded from the analysis due to contamination issues across tissue boundaries.

**Figure 1.**
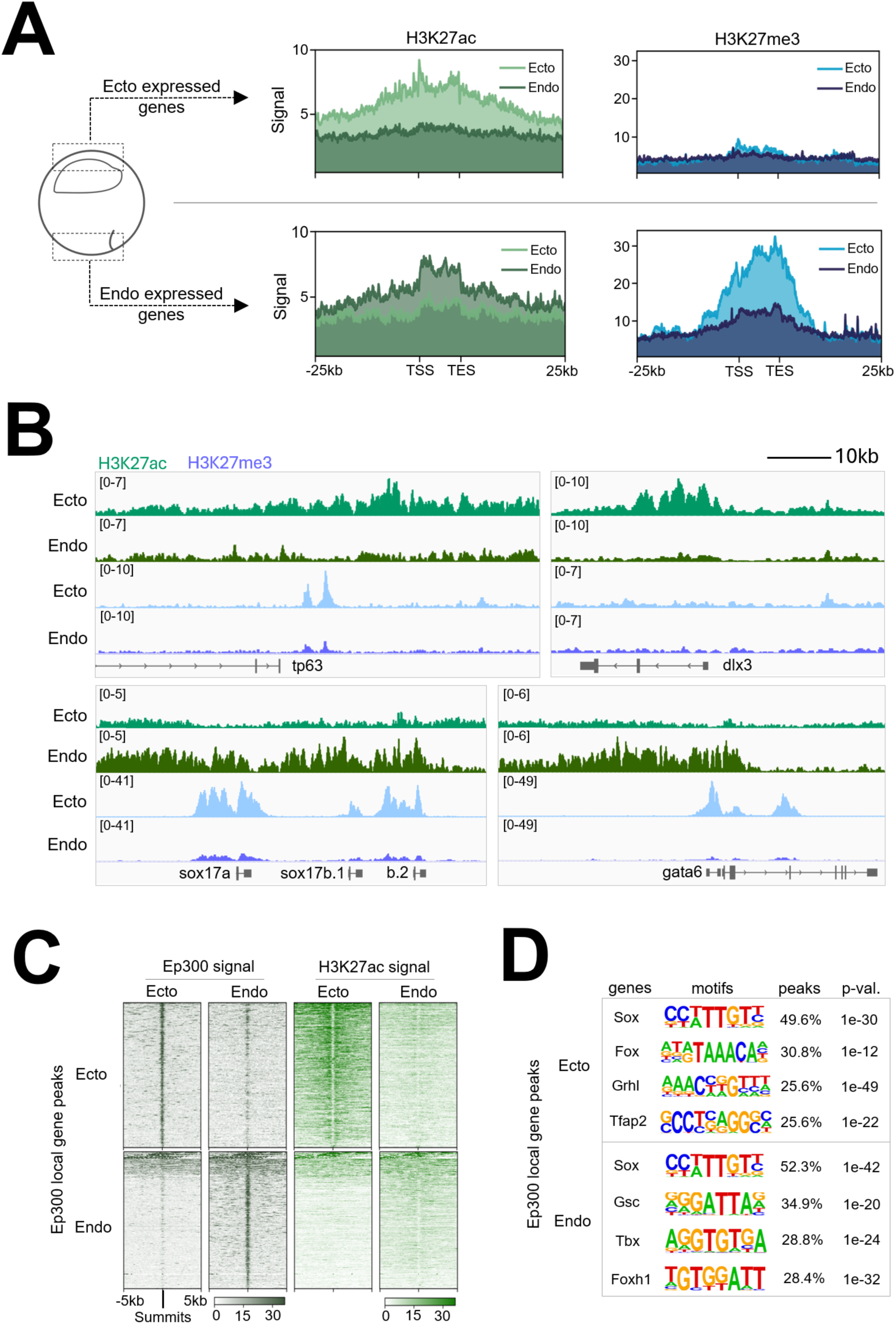
Differential Epigenetic Regulation of Ectodermal and Endodermal Genes in Early Gastrulae. (A) Signal deposition of H3K27ac (left, green) and H3K27me3 (right, blue) across the top 250 zygotically expressed ectodermal (top) or endodermal (bottom) gene regions obtained from early gastrula ectoderm and endoderm dissections. (B) Genome browser tracks showing H3K27ac and H3K27me3 marks along representative zygotically expressed ectodermal and endodermal genes. (C) Signal enrichment of Ep300 (left) and H3K27ac (right) across Ep300 bound regions associated with top 250 zygotically expressed ectodermal (top) and endodermal (bottom) genes from early gastrula ectoderm and endoderm dissections (D) Transcription factor motifs detected within Ep300 peaks within 20kb of top 250 ectodermal (top) and endodermal (bottom) genes.

Ectodermally expressed genes, including both up and downstream regions, are more highly decorated with the activating H3K27ac mark in the ectoderm compared to the endoderm (Figure 1A, B, S1C). On the other hand, endodermally expressed genes are more highly marked by H3K27ac in the endoderm compared to the ectoderm. We also examined the distribution of the repressive mark H3K27me3 across both ectodermally and endodermally expressed genes.

Endodermal genes are strongly marked by H3K27me3 in ectoderm, but relatively remain unmarked by H3K27me3 in endoderm (Fig 1A, B, S1C). This H3K27me3 deposition suggests the mechanism of active repression of endodermal genes in the ectoderm, which prevents their inappropriate expression outside their endodermal environment. In contrast, ectodermal genes lack H3K27me3 enrichment in both endoderm and ectoderm cells. Therefore, in early gastrula embryos, robust accrual of tissue specific “H3K27me3 repressive” marks occur on endodermal genes in the ectoderm, but not ectodermal genes in the endoderm.

Since H3K27ac marking of ectodermal and endodermal genes correlates well with their region-specific expression in developing embryos, we examined the binding of the histone acetyl transferase Ep300 (Figure 1C; S1D), which is a commonly used marker to identify functional enhancers (Heintzman et al., 2009). In ectoderm explants, Ep300 binding is high in ectodermally expressed genes, while in endoderm explants it is preferentially enriched in endodermal gene binding (Figure 1C). Ep300 functions as a transcriptional co-activator and acetyltransferase, but it does not possess a DNA binding domain. Therefore, the recruitment of Ep300 to specific genomic loci requires TF recruitment. To identify potential Ep300 cofactors, TF motif enrichment search of Ep300-bound peaks was performed for both ectodermally and endodermally expressed gene regions. Sox, Fox, Grhl and Tfap2 family motifs were most frequently identified in Ep300 peaks associated with ectodermally expressed genes, whereas the Sox, Gsc, Tbx and Foxh1 motifs were most frequent in endodermally expressed genes (Figure 1D). These results suggest that gene families related to Sox and Fox TFs are critical in both ectodermal and endodermal development. Importantly, *foxi2* and *sox3* transcripts are highly expressed maternally and enriched in the ectoderm (Paraiso et al. 2019; Cha et al. 2012; Zhang et al., 2003; Blitz et al., 2017), implying their involvement, in not only regulating the ectodermal gene regulatory program, but also influencing epigenetic modifications of the ectodermal genes.

### Foxi2 and Sox3 prebind ectodermal CRMs before H3K27ac marks

Previous work showed that maternal Foxi2 can bind to an upstream cis-regulatory module (CRM) in the *foxi1* gene to activate its expression (Cha et al., 2012). We wished to examine the genome-wide regulatory role of Foxi2 during early development. ChIP-seq analysis of Foxi2 was performed in blastula (st. 8, 9) and early gastrula (st. 10.5) embryos and identified a total of 16,427 bound regions over the entire time course. There are 13,158 high confidence Foxi2 binding regions in the mid blastula, 6,668 in the late blastula, and 6,039 in the early gastrula (Figure 2A). Overall, Foxi2 binding is dynamic, displaying unique (class I, III, VI), shared (class II, IV, V), as well as persistently occupied (class VII) binding patterns across stages (Figure 2A, B, Table S2).

**Figure 2.**
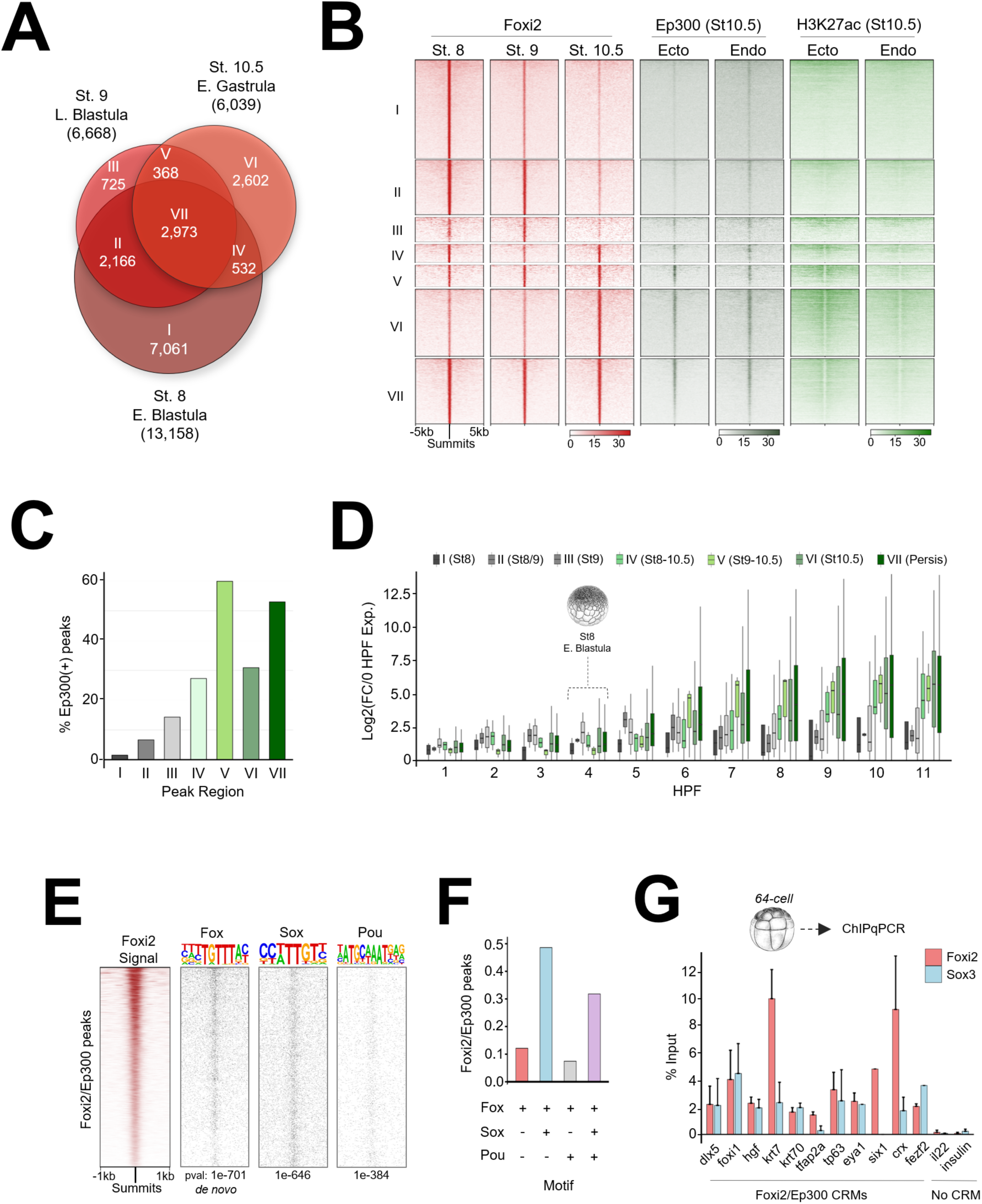
Temporal Dynamics and Motif Analysis of Foxi2 DNA Binding and Co-Occupancy with Ep300 in Early Gastrula. (A) Venn diagram showing temporal dynamics of Foxi2 DNA binding, with ChIP-seq peaks categorized into Classes I-VII. (B) Analysis comparing Ep300 and H3K27ac signal from early gastrula ectoderm and endoderm dissection, within Foxi2 peak classification. (C) Proportion of Foxi2 class I-VII peaks overlapping Ep300 peaks from early gastrula ectoderm. (D) Temporal RNA expression profiles of genes within 20kb of Foxi2/Ep300 co-bound peaks in classes I-VII. (E) Top predicted Fox motif and best matched Sox and Pou motifs identified within Foxi2/Ep300 peaks. (F) Proportion of Fox, Sox and Pou motifs present in Foxi2/Ep300 peaks. (G) ChIP-qPCR analysis of Foxi2 and Sox3 binding at Foxi2/Ep300 co-bound regions from 64-cell embryo chromatin.

To characterize functionally relevant Foxi2 peaks, we examined the correlation of classified Foxi2 peaks with Ep300 and H3K27ac peaks in ectoderm and endoderm explants. Class IV, V, VI and VII regions contain more H3K27ac deposition and Ep300 binding compared to classes I, II and III (Figure 2B, S2A). In total, 3,050 Foxi2 regions are Ep300 co-occupied, where class IV, V, VI and VII Foxi2 peaks show the highest degree of overlap with both Ep300 and H3K27ac (Figure 2B, C, S2A). We bioinformatically assigned nearest zygotic genes to the Foxi2 peaks marked by Ep300, and examined whether these CRMs are linked to transcriptional activity of the genes. We find that class IV, V, VI and VII genes become transcriptionally active during gastrulation, whereas class I, II and III genes remain inactive (Figure 2D). These results suggest that Foxi2 peaks persisting from stage 8-10, are shared from stage 9,10.5, or are specifically bound at stage 10.5 are associated with Ep300 and thus represent active CRMs.

TFs form complexes on CRMs and work together in combination to regulate gene expression. To predict potential co-factors that may work together with Foxi2, we characterized TF motifs enriched within Foxi2 bound regions. As expected, *de novo* motif analysis reveals the most centrally enriched are Fox motifs (Figure 2E). Besides the Fox motif, the next most abundantly present and centrally enriched motifs are Sox and Pou. Among Foxi2 peaks bound by Ep300, ∼78% contain combinatorial Fox/Sox motifs (Figure 2E, F). The Pou motif was also detected, but less frequent than the Fox and Sox motifs (Figure 2E, F). Given that Foxi2 and Sox3 are maternally expressed, we assessed whether Foxi2 and Sox3 prebind functional ectoderm CRMs, before Ep300 binding and the onset of ZGA. Foxi2 and Sox3 ChIP-qPCR analysis of 64-cell stage embryos (st. 6.5, ∼3hpf) reveals that Foxi2 and Sox3 occupy CRMs for *dlx5, foxi1, hgf, krt7, krt70, tfap2a, and tp63* (Figure 2G), which are expressed in the ectoderm of the early gastrula. This demonstrates that Foxi2 and Sox3 prebind to these ectodermal CRMs during cleavage stages, thus prior to ZGA (∼4-4.5hpf), and before the recruitment of Ep300 and the deposition of H3K27ac. Additionally, we discovered that these maternal TFs premark CRMs of *eya1*, *six1, crx,* and *fezf2*, which are expressed in the preplacodal regions (PPR) or anterior neural plate regions of neurula stage embryos (Moody et al., 2015; Maharana and Schlosser, 2018). This finding demonstrates that these maternal TFs recognize the CRMs of future ectodermally expressed genes during the cleavage stages.

### Active ectodermal CRMs are co-occupied by Foxi2 and Sox3

Our ChIP-seq analysis of Sox3 reveals a total of 18,965 peaks, representing 5,828, 4,402 and 14,648 peaks in stage 8, 9, and 10.5 embryos (Figure 3A). In line with our Foxi2 ChIP-seq analyses, Sox3 peaks display temporally unique (class I, III, VII), shared (class II, IV, VI), and persistently occupied (class V) binding patterns (Figure 3A, B). Class V, VI, and VII peaks all displayed higher Ep300 peaks and H3K27ac deposition compared to class I, II, III, and IV (Figure 3B, C), suggesting that class V, VI and VII associated genes are active. These results imply that, like Foxi2, Sox3 peaks that persist from stage 8-10.5 and 9-10.5, or are specifically bound at stage 10.5 are associated with Ep300 and represent active CRMs. We examined whether these CRMs are linked to transcriptional activity of the genes. Class V, VI and VII associated genes are expressed at higher levels compared to class I genes (Figure 3D). Class II, III and IV gene lists were excluded from this analysis due to the low number of uniquely (class specific) bound genes within these categories.

**Figure 3.**
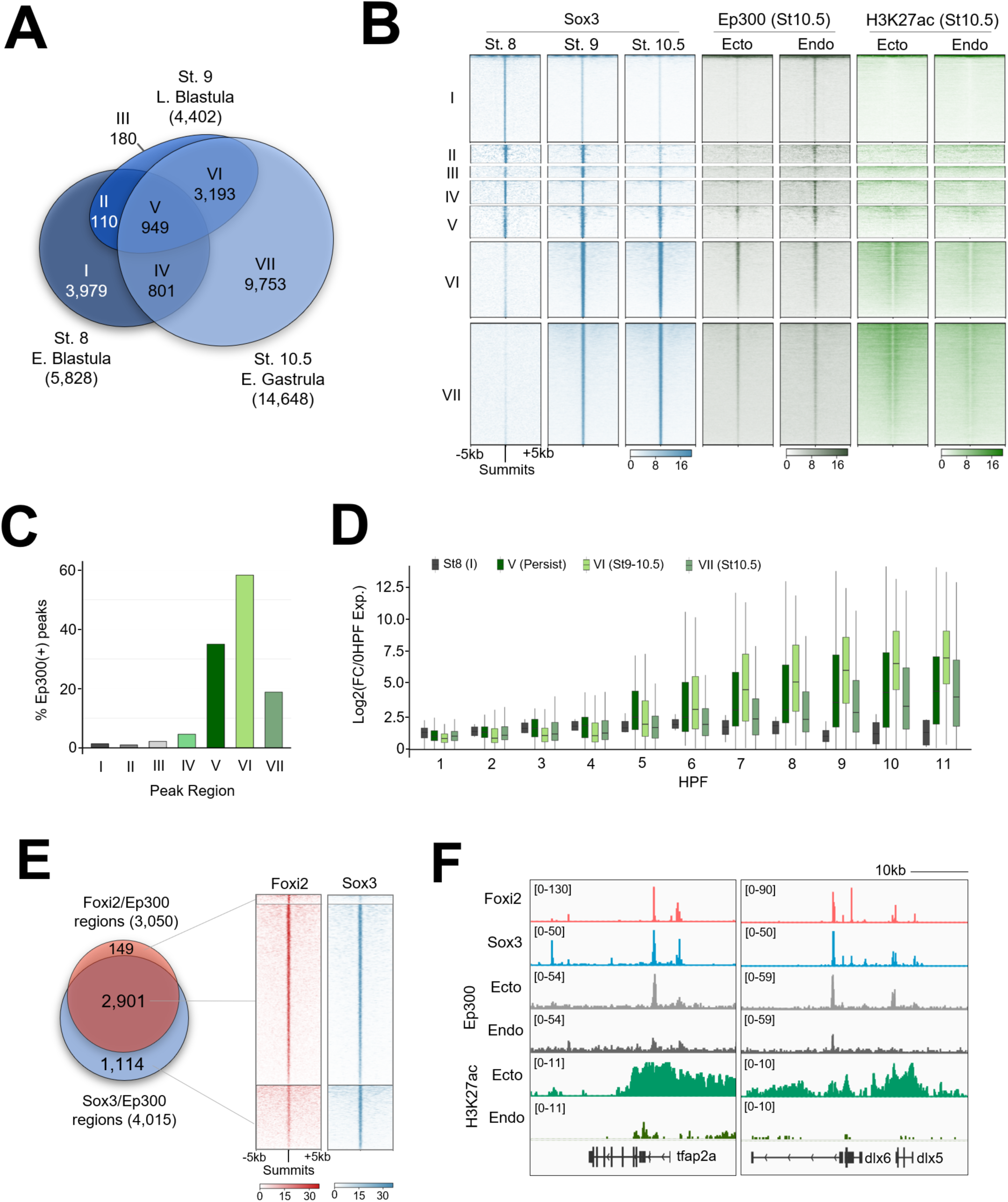
Foxi2 and Sox3 Preferentially Colocalize at Ectoderm CRMs in the Presence of Ep300. (A) Venn diagram illustrating the temporal dynamics of Sox3 ChIP-seq peaks, classified into categories I-VII. (B) Clustering analysis comparing Sox3 peak classifications with Ep300 and H3K27ac signals from early gastrula ectoderm and endoderm dissection. (C) Proportion of Sox3 class I-VII peaks overlapping with Ep300 peaks. (D) Temporal gene expression profiles of genes located within 20kb of Sox3/Ep300 co-bound peaks in Class I-VII. (E) Comparison of Foxi2 and Sox3 co-bound and independently bound regions overlapping with Ep300. (F) Genome browser tracks highlighting selected ectodermally-expressed genes with Foxi2, Sox3 and Ep300 binding.

Next, we examined the genomic colocalization of Foxi2 and Sox3. Examining the binding profile of either TF centered across each other’s peaks reveals their high degree of binding overlap. Among the 4,866 Foxi2 and Sox3 co-bound regions (Figure S3A-C), 2,901 also overlap with ectodermal Ep300 peaks (Figure 3B, C). These overlapping regions account for 95% of the 3,050 Foxi2/Ep300 peaks and 72% of the 4,015 Sox3/Ep300 peaks (Figure 3E). We examined the dynamic binding of Foxi2 and Sox3 around several ectodermal genes and observed that their co-occupancy is either maintained or increases from the early blastula to the early gastrula (Figure S3D). Additionally, these CRM regions show significantly higher Ep300 binding and H3K27ac mark deposition in the ectoderm compared to endoderm, suggesting their functional importance in ectodermal gene regulation. Linking 2,901 Foxi2/Sox3 CRMs co-bound with Ep300 to their nearest genes (Table S3), we find highly significant GO terms related to cell fate specification of neural (e.g., *fezf2, neurog3*, *zic3*), and epidermal lineages (e.g., *dlx3*, *grhl2*, *krt7*) (Figure S3E). This suggests that Foxi2 and Sox3 orchestrate a cis-regulatory code critical for driving both neural and non-neural ectoderm specification.

In early gastrula stage embryos, ectodermal genes such as *dlx5*, *foxi1*, *hgf*, *krt7*, *krt70*, *tfap2a*, and *tp63*, are bound by both Foxi2 and Sox3, and overlap with Ep300 peaks (Figure 3F, S3D). Interesting, these CRMs are also premarked by Foxi2 and Sox3 by the 64-cell stage (Figure 2G). Given the established role of the Foxi family TFs in sensory placode formation (Solomon et al., 2003; Matsuo-Takasaki et al., 2005), we examined the key sensory placode genes, such as *eya1*, *six1*, and the anterior neural marker *fezf2,* which we found to be marked during cleavage stage (Figure 2G). Indeed, these genes are also co-bound by Foxi2 and Sox3 in the early gastrula stage embryos (Figure S3D), sharing the same CRM-pre-bound by Fooxi2 and Sox3 during cleavage stages (Figure 2G). These findings underscore the master regulatory role of maternal TFs in pre-marking critical ectodermal CRMs for future gene expression.

### Independent and cooperative roles of Foxi2 and Sox3 regulate ectodermal gene expression

To investigate the molecular function of Foxi2 and Sox3, we generated Foxi2 and Sox3 knockdown phenotypes using translation-blocking morpholino antisense oligonucleotides (MOs) (Figure S4A, B. Embryos injected with Foxi2 or Sox3 MOs showed substantial depletion of Foxi2 and Sox3 proteins (Figure 4A). Sox3 morphants exhibited decreases in body length and inhibited blastopore closure, while Foxi2 knockdown embryos showed robust defects in both dorsoventral patterning and body length, as well as reduced head size (Figure 4B). RNA-seq analysis of both Foxi2 and Sox3 morphants compared to wild-type embryos identified genes activated and repressed by Foxi2 and Sox3 (Figure 4C, D). We intersected these data with Foxi2 and Sox3 ChIP-seq datasets to identify direct target genes, defined as those showing at least a 2-fold change in expression in the morphants and having Foxi2 and/or Sox3 binding sites within 20kb of the gene (Figure 4C, D, Table S4). At the late blastula stage, Foxi2 and Sox3 directly activate 78 and 114 genes, and repress 68 and 18 genes, respectively (Figure 4C, Table S4). A similar analysis in gastrula-stage embryos revealed that Foxi2 and Sox3 directly activate 72 and 164 genes and repress 54 and 136 genes, respectively (Figure 4D, Table S4). Intersecting the lists of genes activated by Foxi2 and Sox3 identifies 35 genes co-bound and co-regulated by both factors. These genes include well-studied ectodermal transcription factors such as *dlx3, dlx5, foxi1, hes5.1, tfap2a,* and *tp63* (Table S4) (Moody and LaMantia, 2015; Maharana and Schlosser, 2018). RT-qPCR analysis confirms that these ectodermal TFs are jointly regulated by both factors (Figure 4E). Separately, 115 genes are regulated solely by Foxi2, while 243 genes are regulated solely by Sox3. Specific targets of Foxi2 include *amotl1, gata3, kit,* and *krt70*, while direct targets of Sox3 include *cdx4, efna3, foxb1, foxd4l1.2, neurog3, olig4, sox21, zic3,* and *zic4.* These data suggest that Sox3 and Foxi2 regulate their specific target genes both jointly and independently, serving as master TFs that govern the early ectodermal program.

**Figure 4.**
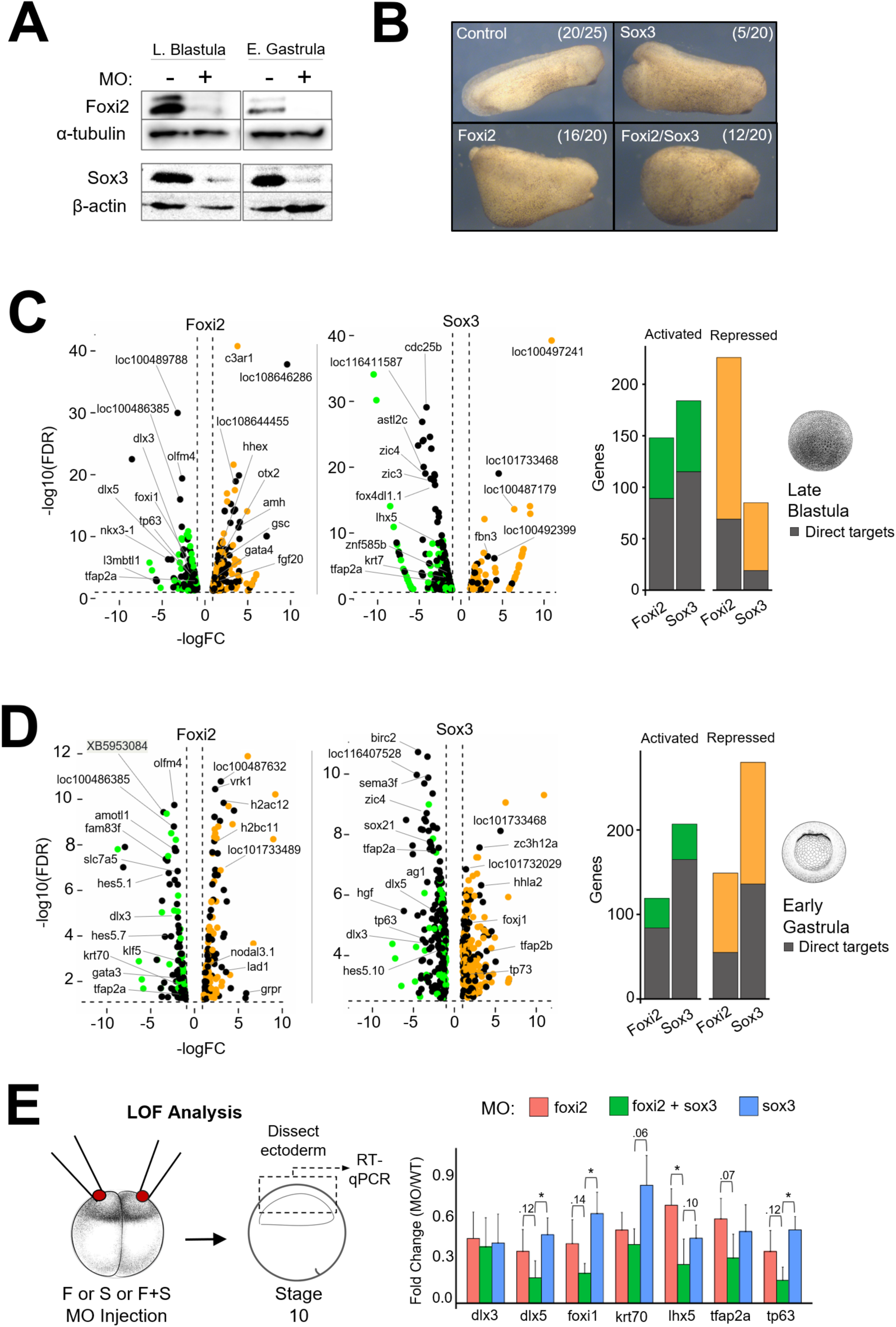
Independent and Cooperative Roles of Foxi2 and Sox3 in Regulating Ectodermal Gene Expression. (A) Western blot of Foxi2 and Sox3 morphant embryos at late blastula and early gastrula stages. (B) Phenotypic analysis of Foxi2, Sox3 and Foxi2/Sox3 morphants at the early tailbud stage. (C, D) RNA-seq analysis of late blastula embryos (top) and early gastrula embryos (bottom) showing activated and repressed genes within 20kb of Foxi2/Sox3 bound sites. Histograms show the total number of direct target genes affected in morphants. (E) Schematic diagram of the experiment (left). RT-qPCR analysis of key ectodermal genes (right) in early gastrula ectoderm of Foxi2, Sox3 and Foxi2/Sox3 morphants. Asterisks indicate statistically significant differences (p< 0.05).

### Foxi2 and Sox3 directly co-regulate spatially distinct inner and outer layer cell states

To gain a deeper understanding of gastrula ectoderm differentiation than can be provided by bulk RNA-seq analyses, we assessed nascent transcripts at the single cell level by performing single nucleus (sn) RNA-seq on stage 10.5 wild-type embryos. We then characterized the cell-type specific expression of Foxi2 and Sox3 direct target genes within this dataset. Pre-processing using a 1500 genes/cell cut-off yielded 13,711 high quality early gastrula nuclei. *De novo* clustering and marker gene expression analysis identified a total of 13 defined cell states, consisting of 5 ectodermal, 4 mesodermal, and 4 endodermal cell types (Figure 5A, Figure S5A-G). The five ectodermal cell clusters map at the bottom of two major lobes in the UMAP (Figure 5A) and markers show that these represent inner neural plate (INP), inner neural plate border (INPB), inner non-neural ectoderm (INNE), outer neural ectoderm (ONE), and outer non-neural ectoderm (ONNE). Notably, inner ectoderm cell clusters are positioned in the left lobe, while outer ectoderm cell clusters segregate in the right lobe (Figure 5A, S5C,E). Two small lobes in the upper right (gray) contain unannotated cells (UC) that may represent a novel cell state characterized by *dscaml1* and *tp73* expression (Figure S5D), but the identity of these cells remains unclear at present.

**Figure 5.**
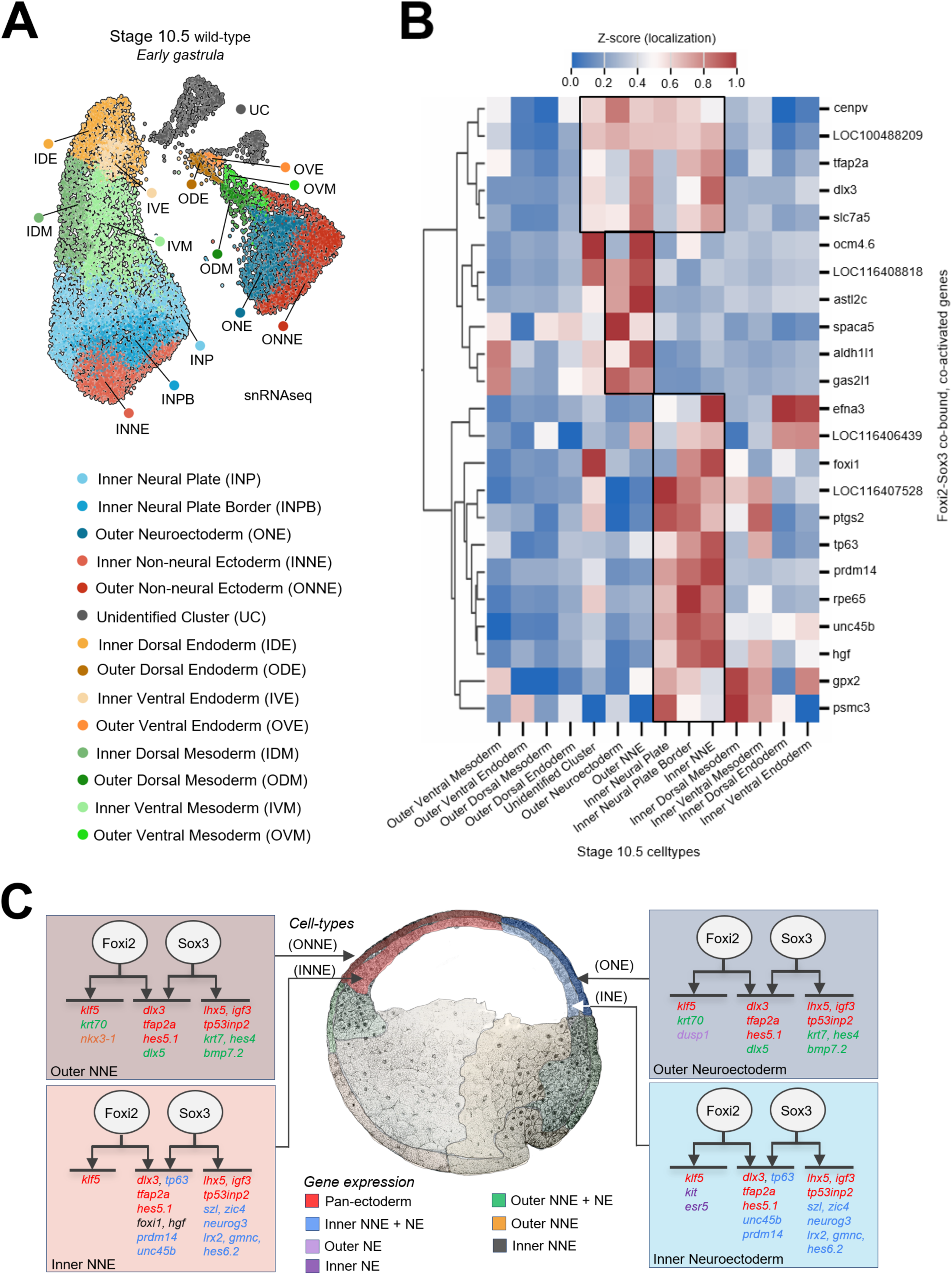
Foxi2 and Sox3 Directly Co-regulate Genes in All Ectodermal Cluster. (A) Single nucleus RNA-seq UMAP (uniform manifold approximation projection) identifying 13 distinct cell types in the early gastrula. (B) Z-score expression analysis of Foxi2/Sox3 co-bound target genes shows distinct gene expression patterns marking outer and inner ectodermal cell types (C) Schematic diagram depicting the regulatory roles of Foxi2 and Sox3, either independently or jointly, in controlling the expression of target genes across various ectodermal cell types.

After identifying 5 ectodermal cell clusters in stage 10.5 embryos, we mapped the expression patterns of Foxi2 and Sox3 direct target genes in the gastrula ectoderm to determine how Foxi2 and Sox3 contribute to the regional identity of ectodermal cells. There are 18 inner layer and 14 outer layer direct ectoderm target genes that are exclusively regulated by Foxi2, which include *amotl1, klf5, krt70, lhx3, nkx3-1 and olfm4* (Table S4). Additionally, 53 inner and 49 outer layer direct ectoderm target genes are exclusively regulated by Sox3, including *foxd4l1.2, neurog3, olig3, olig4, sal3, sox21, szl, zic3, and zic4,* many of which are involved in neurogenesis (Table S4). Pan-ectodermal targets (e.g., *dlx3, hes5.1, klf5, lhx5* and *tfap2a*) are expressed across all five cell states. Some of these genes are specifically regulated by Foxi2 or Sox3 alone, while others require the combinatorial input of both Foxi2 and Sox3 (Figure 5C). Additionally, we identified target genes of Foxi2 and Sox3 that are specifically expressed in the inner (both non-neural and neural) ectodermal layer, such as *foxi1, prdm14, szl, tp63,* and *zic4*. Similarly, genes such as *dlx5, hes4* and *krt70* are expressed in the outer (both non-neural and neural) ectodermal cell states. Since these genes are regulated by Foxi2 and/or Sox3 in these distinct ectodermal layers, we suggest that additional factors, like aPKC, differentially localized or activated in these layers, contribute to the segregation of the inner and outer ectoderm (Chalmers et al. 2003). .

### Foxi2 and Sox3 are master regulators shaping the ectodermal epigenetic landscape

Given the potential synergy of Foxi2 and Sox3 within the ectoderm and their distinct roles in ectoderm target gene activation, could Foxi2 and Sox3 together serve as master regulators of ectodermal differentiation? To test this hypothesis, we injected Foxi2 and/or Sox3 mRNA into the vegetal pole and assessed the endoderm cell fate in dissected vegetal tissue explants at gastrula stage 10.5 (Figure 6A). While overexpression of Foxi2 or Sox3 alone induced modest ectoderm gene expression, coinjection of both factors robustly activated the ectodermal genes, *dlx5*, *foxi1, krt70*, *lhx5* and *tfap2a*. Moreover, expression of key endodermal genes such as *gata4*, *gsc*, *hhex* and *otx2* was significantly reduced in endodermal explants co-expressing Foxi2 and Sox3 mRNA. These findings support Foxi2 and Sox3 working together as master regulators of ectodermal identity.

**Figure 6.**
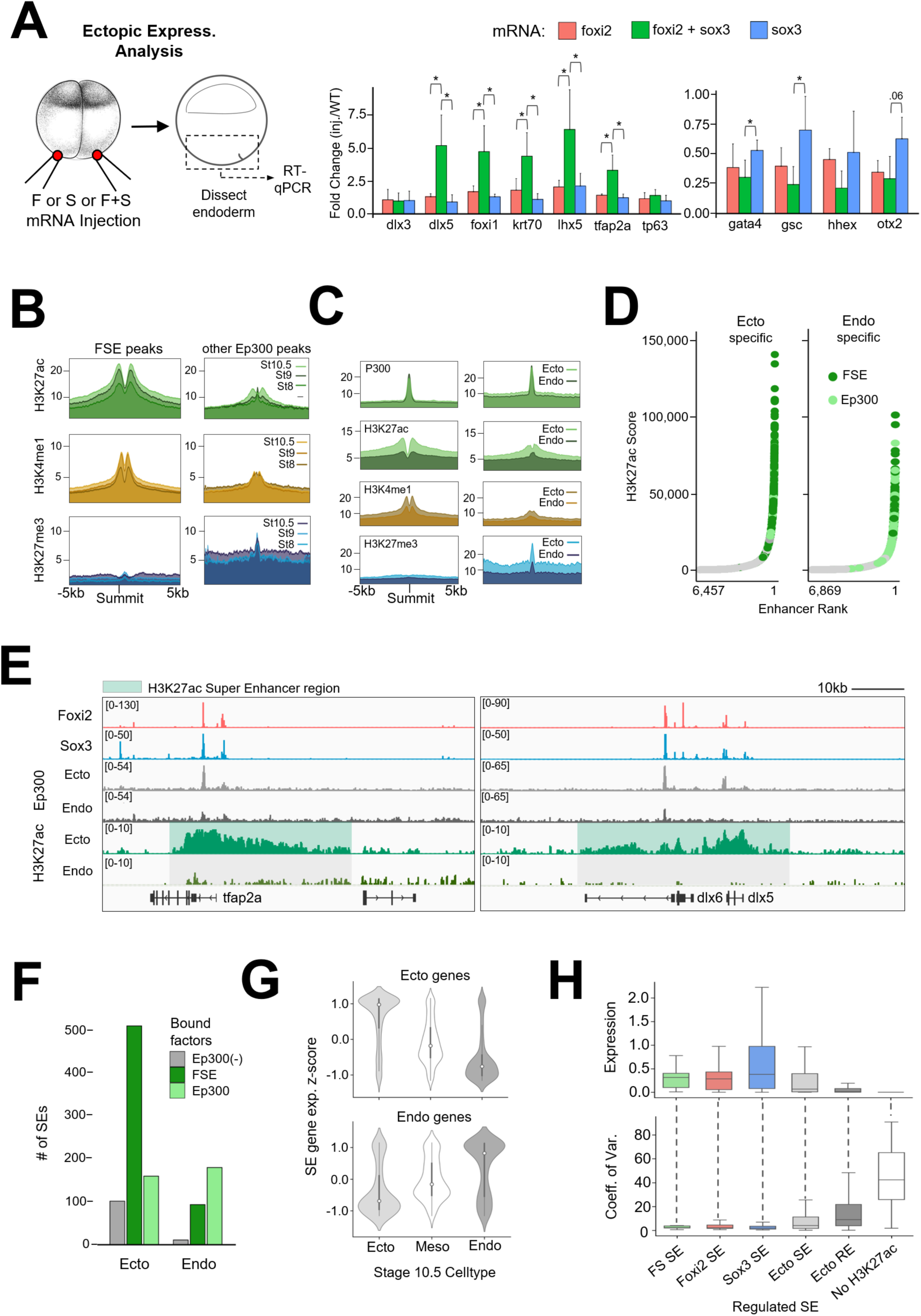
Foxi2 and Sox3 are Master Regulators Shaping the Ectodermal Super Enhancer Landscape. (A) Schematic diagram of the experimental design (left). RT-qPCR analysis of key ectoderm and endoderm genes from early gastrula endoderm after ectopic expression of Foxi2, Sox3 or Foxi2/Sox3 mRNA. Asterisks indicate statistically significant differences (p< 0.05) (B) Accumulation of H3K27ac, H3K4me1 and H3K27me3 histone modifications around Ep300 peaks. (C) Deposition of Ep300, H3K27ac, H3K4me1 and H3K27me3 at Ep300 peaks present in the early gastrula ectoderm and endoderm. (D) Peaks co-occupied by Foxi2/Sox3/Ep300 (FSE) are more frequently associated with ectodermal SEs than with endodermal Ep300-only peaks (Ep300) (E) Genome browser views of ectodermal SEs (green shaded area), which are present in ectoderm but absent in endoderm. (F) Histogram comparing the total number peaks associated with super enhancers containing Foxi2/Sox3 and Ep300 (FSE), Ep300 only (no Foxi2/Sox3 binding), and Foxi2/Sox3 without Ep300 (Ep300(-)). FSE peaks are highly associated with ectodermal SEs. (G) Z-score expression analysis of genes located within 20kb of ectoderm (top) and endoderm (bottom) SEs. snRNA-seq cell-type expression analysis reveals that genes associated with ectodermal SEs are preferentially expressed in ectodermal cells (H) Average expression levels of Foxi2-and/or Sox3-regulated ectodermal SE-associated genes compared to all enhancer associated genes (top). Coefficient of variation (COV) analysis shows that Foxi2 and/or Sox3-regulated ectodermal SE associated genes exhibit less variability in expression compared to all enhancer associated genes using snRNA-seq.

In addition to the transcriptional role of Foxi2 and Sox3 in specifying the ectodermal cell state, we determined whether Foxi2 and Sox3 also influence the epigenetic state of the ectodermal cells. Temporal analysis (Figure 6B) of H3K27ac and H3K4me1 signal deposition around 2901 CRMs that co-bind Foxi2, Sox3, and Ep300 (FSE), (Figure 3E) reveals an increase from the early blastula to the early gastrula stage. Interestingly, the repressive mark, H3K27me3 becomes gradually enriched around Ep300 peaks lacking Foxi2 and Sox3 binding (Figure 6B). These results suggest that Foxi2/Sox3 co-binding promotes H3K27ac and H3K4me1 enrichment at the expense of H3K27me3 deposition around ectodermal CRMs.

To overcome the lack of germ layer resolution in whole-embryo histone marking studies, we also performed experiments using dissected ectodermal and endodermal tissues from the early gastrula stage. While no significant differences in Ep300 binding were observed between ectoderm and endoderm at both Foxi2-Sox3 co-bound (FSE) regions and Ep300 only peaks (lacking Foxi2/Sox3) (Figure 6C), we found a significant difference in histone modifications. In the ectoderm, FSE regions showed significant accumulation of H3K27ac and H3K4me1, while H3K27me3 deposition is notably absent (Figure 6C). In contrast, Ep300-only peaks showed modest H3K27ac and H3K4me1 accumulation but displayed strong H3K27me3 enrichment. These findings suggest that Foxi2 and Sox3 specifically regulate the epigenetic landscape of ectodermal genes within the ectodermal germ layer, but not in the endoderm.

### Foxi2 and Sox3 marked regions establish ectodermal super enhancers to ensure robust gene expression

The high levels of H3K27ac modifications around Foxi2 and Sox3 co-bound sites raises the question of whether some of these regions are located within super enhancers (SEs), which are large clusters of enhancers characterized by high levels of TFs, coactivators (such as Mediator), and chromatin modifiers. SEs are typically identified by analyzing genomic regions highly marked by H3K27ac (Whyte et al., 2013). H3K27ac ChIP-seq analysis of ectodermal and endodermal explants identified 14,074 ectoderm and 15,545 endoderm enhancer peaks (“regular” enhancers, REs), with 9,554 being shared between the two germ layers (Figure S6A). To further investigate, we processed the data to identify SEs in both ectoderm and endoderm based on their increased size and H3K27ac deposition relative to the enhancer landscape (Figure 6D, S6A). The average size of both ectodermal and endodermal SEs are ∼20kb per locus, which is significantly larger than “regular” enhancer (RE) loci, which are typically less than 2kb (Figure S6B). While 403 SE regions overlap between ectodermal and endodermal SEs, 767 are unique to the ectoderm, and 283 are unique to the endoderm (Figure S6A). These findings highlight that ectodermal SEs are more abundant than endodermal SEs and the timing of accumulation of H3K27ac suggests distinct germ layer SEs arise between the onset of ZGA and early gastrulation (Figure 6B).

Highlighting the locations of gene’s associated with FSE-bound regions on a super enhancer ROSE (ranked order of super enhancers) plot revealed that the Foxi2 and Sox3 regulated genes are highly enriched in ectodermal SEs relative to REs (Figure 6D,E,F) or endodermal SEs (Figure S6B,D). The expression levels of ectoderm SE genes associated with Foxi2/Sox3 bound regions containing Ep300 (FSE) are also significantly higher than SE genes with no Ep300, and RE-associated genes (Figure S6E) during embryonic development. Using our snRNA-seq data (Figure 5), we found that ectodermal SE-associated genes are preferentially expressed in the ectodermal germ layer (Figure 6G), and also in ectodermal sub-lineages (Figure S6F). In contrast, endodermal SE-associated genes are preferentially expressed in endodermal germ layer and its endodermal sub-lineages.

Finally, we investigated the effect of SEs on gene expression variance (transcriptional “noise”) across individual cells. To quantify differences in gene expression across genes, we measured the coefficient of variation (COV), which reflects the extent of gene expression among individual cells. Analysis of snRNA-seq data revealed that ectodermal SE-associated genes show higher gene expression levels and a lower COV, compared to RE genes and genes lacking H3K27ac deposition (Figure S6G). Additionally, we observed that SE-associated Foxi2/Sox3, Foxi2-only, and Sox3-only regulated target genes display the highest expression, and the lowest COV, compared to the entire landscape of ectoderm SEs and ectoderm REs and genes unassociated with an enhancer (Figure 6H). These results suggest that ectoderm SEs, enriched with Foxi2 and Sox3 binding, drive the most robust and stable expression of ectoderm-specific genes, while minimizing transcriptional noise. In sum, at the onset of gastrulation Foxi2 and Sox3-marked regions preferentially acquire Ep300 and accumulate H3K27ac to establish SEs, which ensure robust gene expression across all ectoderm sub-lineages.

## DISCUSSION

In this study, we identified maternal Foxi2 and Sox3 as master regulators of the ectodermal germ layer in *Xenopus*, based on the following evidence: (1) ectopic expression of these TFs in endodermal cells confers ectodermal cell fate specification; (2) loss-of-function analysis results in the loss of key ectodermal markers; (3) bound sites for these TFs correlate with Ep300 recruitment and the marking of ectodermal CRMs with H3K27ac; and (4) the co-bound regions include SEs that drive a robust ectodermal gene expression program. Since Foxi2 and Sox3 prebind to many CRMs that are critical for the later activation of ectodermal developmental programs (e.g., sensory placode formation), we propose a model in which animally enriched Foxi2 and Sox3 establish early genomic occupancy at these CRMs during the cleavage stage. These interactions with the genome prefigure the accumulation of activating epigenetic modifications around these elements, by early gastrulation, driving the formation SEs and spatially distinct gene regulatory programs that define ectodermal identity.

### The roles of maternal Foxi2 and Sox3 in early development

In vertebrates, the Foxi subfamily comprises Foxi1, Foxi2, and Foxi3, which are pivotal in epithelial differentiation and organogenesis (Edlund et al., 2015). There are limited studies performed on the function of *foxi2* in early development. However, the available data consistently show that *foxi2* has roles in later stages of development, such as establishment of sensory placodes and anterior neural ectoderm. In the chick embryo, Foxi2 is crucial for craniofacial development, regulating the formation of the pharyngeal arches (Khatri and Groves, 2013). In mouse, Foxi2 is expressed in the developing forebrain, neural retina, dental and olfactory epithelium (Hulander et al., 1998, Ohyama et al., 2004, Wijchers et al., 2006). And in *Xenopus*, Foxi2 morphants similarly appear to have anterior structure defects (Figure 4B).

Despite these conserved roles of Foxi2 in postgastrulation vertebrate embryos, evidence for maternally expressed Foxi2 being essential for early ectodermal germ layer cell lineage development is currently limited to *Xenopus.* Maternally expressed Foxi2 directly activates *foxi1* (Cha et al., 2012, Figure 4C,E, S4A), which is best characterized for its key roles in mucociliary development, ionocyte specification, and sensory placode formation in *Xenopus* (Bowden et al., 2024). This is consistent with the established roles of Foxi1 in other vertebrates. Interestingly, *Xenopus* Foxi2 pre-binds to the CRMs of *foxi1* as well as sensory placode genes such as *eya1*, *six1,* and *fezf2* during cleavage stages (Figure 2G), perhaps priming these CRMs for subsequent activation during later stages of development.

Based on the evidence, we speculate that all three *Xenopus foxi* genes are involved in ectodermal specification, but their roles may have diverged during evolution. The maternal function of *foxi2* in early ectoderm specification may not be unique to early *Xenopus* embryos and might exist in other species but hasn’t yet been studied. In axolotl embryos, RNA-seq analysis shows that *foxi2* mRNA is detectable before zygotic transcription, suggesting it is maternally expressed in this caudate amphibian (Jiang et al., 2017). A zebrafish *foxi* gene referred to as *foxi1*, but which our synteny analysis (unpublished observations) suggests is orthologous to *Xenopus* and mammalian *foxi2*, is first expressed shortly after the onset of ZGA as indicated by RNA-seq profiling (White et al., 2017; https://www.ebi.ac.uk/gxa/home). This suggests it might be more broadly expressed and have an earlier role in development in teleost fish. Finally, a lamprey *foxi* gene that is syntenic to both *foxi1* and *foxi2* (our unpublished observations) is also expressed broadly in the animal pole at blastula stage (York et al., 2024). This expression pattern suggests activity in the early ectoderm, implying conserved functions that predate the divergence of fishes (York et al., 2024). In *Xenopus*, the *foxi1* and *foxi3* genes are not maternally expressed, but are transcribed after onset of ZGA. *foxi1* and *foxi3.2* (but not *foxi3.1*) are expressed broadly in the blastula ectoderm and later become confined to epidermal lineages, while being excluded from the neural plate (Lef et al., 1994; Pohl et al., 2002; Matsuo-Takasaki et al., 2005). Overexpression and loss-of-function experiments of various kinds suggest that these genes might have either redundant functions to *foxi2,* or more likely act as downstream effectors induced by *foxi2*. This is particularly the case with *foxi1.* Therefore we hypothesize that the early role(s) of *foxi2* in ectodermal germ layer specification, while being to induce numerous genes, includes activation of *foxi1* and *foxi3.2*. Similar effects on ectodermal specification in other anamniotes might involve these factors and zygotically expressed *foxi2,* but further experiments in these species are needed to confirm the model.

In *Xenopus*, *sox3* expression is both spatially and temporally dynamic during development. *sox3* mRNA is maternally supplied with expression throughout the ectoderm prior to gastrulation and becomes restricted to the presumptive neural plate by mid-gastrula (Zhang et al., 2003; Penzel et al., 1997). Sox3 represses endoderm formation to facilitate proper germ layer formation (Zhang et al., 2003, Zhang and Klymkowsky, 2007). In later stages, *sox3* is expressed in neural progenitors along the neural tube, where it is implicated in promoting neural identity and preventing premature differentiation. Recently, Sox3 has been shown to have a role in ectoderm pluripotency, and also function as a pioneer TF affecting chromatin accessibility (Buitrago-Delgado et al., 2018; Gentsch et al., 2019). Our findings suggest that Sox3 has an additional role in orchestrating the ectodermal program during germ layer formation. This is supported by evidence showing that 1) several critical ectodermal genes are direct targets of Sox3, 2) *sox3* knockdown results in down-regulation of ectodermal gene expression, and 3) ectopic expression of Sox3 and Foxi2 upregulates ectodermal gene expression in the endoderm while repressing mesendodermal markers. it is plausible that Foxi2 primes pre-bound CRMs during early development and subsequently hands off its regulatory roles to other factors, such as Foxi1 or Foxi3.2, which may be induced by Foxi2. This handoff mechanism could maintain CRM accessibility by preventing silencing through repressive histone modifications, thereby ensuring readiness for transcriptional activation in later developmental stages.

### Advantages of germ layer specific SE formation in rapidly dividing embryos

We identified germ layer-specific SEs based on H3K27ac accumulation across the *Xenopus* embryonic genome. Notably, ectodermal SEs are highly correlated with the binding of Foxi2 and Sox3 TFs (Figure 6D). Our findings also show that SE-associated Foxi2/Sox3 target genes exhibit higher expression levels compared to genes associated with REs, while displaying reduced expression noise (Figure 6H). Based on these germ layer specific SE activities, we propose that SE-linked gene expression contributes to developmental stability during cell fate specification, thereby enhancing the robustness of embryonic tissue formation.

Rapidly dividing embryos, such as those of *Xenopus* and zebrafish, undergo swift transitions from gastrulation to neurulation within a short developmental timeframe. These aquatic embryos also face external influences, such as temperature, that impact their development. To ensure efficient and error-free progression, these embryos must not only achieve a rapid surge in gene expression during early stages but also tightly regulate expression to avoid critical errors.

Super-enhancers (SEs) offer a solution by concentrating transcription factors (TFs) and cofactors at critical genomic regions, enabling robust and precise gene expression. We propose that in these rapidly dividing embryos, a key strategy might involve pre-loading enhancers with TFs before zygotic genome activation (ZGA). While direct evidence from ChIP-seq at cleavage stages is lacking, limited ChIP-qPCR data suggest that pre-bound TFs may prime enhancers for rapid and robust activation at ZGA. At the onset of ZGA, recruitment of Ep300, Kmt2c/d, and RNA polymerase II to gene bodies may enable efficient transcription, ensuring developmental success in these time-constrained systems.

### Factors involved in segregating ectodermal cell clusters

The superficial (outer) and inner layer cells in *Xenopus* embryos have distinct transcriptomes in our snRNA-seq data (Figures 5 and S5). These differences arise due to distinct cues that are present by the 64- to 128-cell stages (Chalmers et al., 2003). The localized expression of *tp63* in the inner (basal) layer and *krt70* and *hes5.10* in the outer layer has been detected through WMISH (Chalmers et al., 2002; 2006). Additionally, our snRNA-seq analysis revealed enrichment of new markers, such as *smad6, col18a1* and zeb in the inner layer, *epas1, nectin2 and atp1b2* in the outer layer, among others (Table S5) expanding our understanding of ectodermal lineage-specific gene expression patterns.

By examining the expression patters of Foxi2 and Sox3 direct target genes in our UMAP of the early gastrula embryo (Figure 5), we discovered that these TFs coordinate the expression of pan-ectodermal genes. For example, genes such as *dlx3 and tfap2a* are co-regulated by both factors, while *klf5* expression is mainly regulated by Foxi2, and *lhx5, igf3* and *tp53inp2* are regulated by Sox3. Since all these genes are expressed across the five ectodermal cell clusters (Figure 5B, C, Table S4), we propose that Foxi2 and Sox3 exhibit both overlapping functions and distinct, nonoverlapping roles in ectoderm development. Among Foxi2 and Sox3 targets, genes like krt7, krt70, dlx5, and hes5/10 are exclusively expressed in the outer ectoderm, while prdm14, unc45b, szl, zic4, and neurog3 are specific to the inner ectoderm. This differential expression suggests that additional factors uniquely active or expressed in each layer may play a role. This highlights that other regulatory mechanisms are superimposed on Foxi2/Sox3 regulated targets that underlie this regionalization of ectodermal cells occurring early in gastrulation.

Two notable candidates involved in this process are the kinases Prkci (also known as aPKC) and Mark3. Prkci is localized in apical cell membranes by the 4-cell stage (Chalmers et al., 2003) and contributes to the segregation of outer (superficial) ectodermal cells at later stages (Ossipova et al., 2007). On the other hand, Mark3 is localized basally and influences the inner layer of cells (Ossipova et al., 2007). This interplay may be essential for initiating outer and inner ectodermal differences that intersect with Foxi2/Sox3 regulated gene expression in the early *Xenopu*s embryo.

Additional factors potentially involved in outer and inner ectoderm development include the Grainyhead-like (Grhl) TFs. Our motif analysis in regions bound by Ep300 revealed a significant enrichment of the Grhl motif in early gastrula ectoderm, but not in endoderm (Figure 1D). The *Grhl* gene family is comprised of three highly conserved members in vertebrates (*Grhl1-3*) (Kudryavtseva et al., 2003; Ting et al., 2003b; Wilanowski et al., 2002). Among these, *grhl1* is the only maternally expressed family member in *Xenopus*, and also continues to be expressed zygotically in both the outer and inner layer of the ectoderm based on our snRNA-seq data. Disruption of *grhl1* activity in *Xenopus* results in severe defects in epidermal differentiation, directly affecting keratin gene expression through binding of Grhl1 to the promoter region of *krt12.4* (Tao et al., 2005). Similarly, *Grhl1* knockout models in both mouse and zebrafish exhibit disrupted epidermal cell differentiation (Wilanowski et al., 2008; Janicke et al., 2010). Both *grhl2* and *3* are zygotically expressed in *Xenopus,* starting during gastrulation and continuing into later stages (Chalmers et al., 2006; Owens et al., 2016). *grhl2* is expressed in the inner layer while *grhl3* is expressed in the outer layer of gastrula ectoderm and later in the superficial layer of epidermis (Chalmers et al., 2006). In mice, loss of *Grhl2* causes non-neural ectoderm disruption, leading to disorganized cell junctions, aberrant mesenchymal protein vimentin, and decreased epithelial integrity (Ray and Niswander, 2016). *Grhl3-*deficient mouse embryos die shortly after birth due to impaired skin barrier function (Ting et al., 2005). We observed that a Grhl motif is not enriched in Foxi2- and Sox3-bound regions (Figure 1D), suggesting that *grhl1* likely regulates a distinct set of ectodermal target genes compared to those of Foxi2 and Sox3. We propose that Grhl1 functions as a maternal ectodermal TF that alongside work with Foxi2 and Sox3 to initiate the global ectodermal specification programming, potentially through a separate pathway. Additionally, *grhl2* and *grhl3* may take over zygotic functions of *grhl1*, operating within the two ectodermal layers during later stages of development. Our future goals focus on dissecting the regulatory pathways of the ectoderm by combining loss-of-function analysis of maternal TFs with single-nucleus transcriptomics. This approach aims to determine the timing of lineage segregation and uncover the contributions of the maternal TFs to the specification of various ectodermal cell lineages and epigenetic changes.

### Experimental Procedures

#### Chromatin Immunoprecipitation (ChIP) Assays

Antibodies for Foxi2 (Cha et al., 2012) and Sox3 (Zhang et al., 2004) were validated previously, while Ep300 antibody was Santa Cruz sc-585. *X. tropicalis* ChIP-qPCR was performed as described in Chiu et al. (2014), using whole embryos or dissected tissue fragments with specific primer sets (Table S6). ChIP-seq libraries were generated using NEB E7645S DNA sequencing kit or the Bioo Scientific NEXTflex ChIP-seq kit, verified using an Agilent Bioanalyzer 2100, and sequenced at UC Irvine’s genomics core facility using the Illumina Novaseq platform.

#### Western Blot

Embryos were homogenized in 1xRIPA buffer (50mM Tris-HCl pH7.6, 1% NP40, 0.25% Na-deoxycholate, 150mM NaCl, 1mM EDTA, 0.1% SDS, 0.5mM DTT) supplemented with Roche’s cOmplete protease inhibitor. The homogenate was centrifuged at 14,000 rpm, and the supernatant was collected and centrifuged again. Western blotting was performed on the final supernatant using anti-Foxi2, anti-Sox3, alpha-tubulin (Sigma, T6168), or beta-actin antibodies (Sigma, A5316).

#### RNA Assays

Total RNA was extracted from embryos using TRIzol reagent (ThermoFisher). PolyA-selected mRNA transcripts were isolated using NEBNext PolyA mRNA Magnetic Isolation Module (NEB E7490S). RNA libraries were prepared using NEBNext Ultra II Library Preparation Kit (NEB E7770S), validated using an Agilent Bioanalyzer 2100, and sequenced on the Illumina Novaseq platform. RT-qPCR analyses were performed using primers listed in Table S6. Reverse transcription was performed using the MMLV reverse transcriptase (ThermoFisher Superscript II). qPCR was carried out on a Roche Lightcycler 480 II using Roche SYBR green I master mix.

#### Morpholino Knockdown, Rescue and Vegetal RNA Injections

*foxi2* translation blocking morpholino was created by GeneTools, Inc. 5’-TTATGAAGTCTGGTGGGACATTCAC-’3. Morpholino rescue was performed using mRNA prepared from a pCS2+ FLAG-*foxi2* construct. *sox3* translation blocking morpholino is 5‘-GTCTGTGTCCAACATGCTATACATC-3’. *sox3* morpholino rescue was performed using a pCS2+ FLAG-*sox3* construct. The *X. tropicalis foxi2* coding sequence was acquired from the *X. tropicalis* Unigene library (TGas144f13) in the pCS107 vector and the *X. tropicalis sox3* coding sequence was cloned into the pCS2+ vector. The *foxi2* plasmid was linearized with HpaI, and the *sox3* plasmid with XbaI, for generating capped mRNA using the SP6 mMessage Machine kit. Morpholino knockdowns were performed by injecting directly into opposing sides of the animal cap at the 1-cell stage or into each animal blastomere at the 2-cell stage. Animal caps were dissected one hour prior to harvesting for RNA preparation. Vegetal RNA injections were performed using indicated amounts of mRNA per embryo into opposing sides of the vegetal mass at the 1-cell stage, or in each vegetal blastomere at the 2-cell stage. Vegetal masses were dissected at blastula stage ∼1 hour before harvesting with Trizol.

#### Nuclei Preparation and snRNA Sequencing

Nuclei were isolated from stage 10.5 embryos using previously published discontinuous sucrose gradient protocols followed by a separate centrifugation through 80% glycerol (Wormington and Brown, 1983; Wolffe, 1989; Nakayama et al., 2022). Briefly, 200 embryos were homogenized in a 250mM sucrose solution containing glycerol and snap frozen in liquid nitrogen (Nakayama et al., 2022). Homogenates were later thawed on ice, brought to ∼2.2M sucrose and centrifuged through a 2.4M sucrose layer at 130,000g for 2 hours at 4oC using a Beckman SW55Ti rotor, followed by a 3400xg centrifugation for 10 min at 4oC through an 80% glycerol cushion. RNAse (NEB) and protease inhibitors (Roche 7x cOmplete, Mini, EDTA-free) at 0.2 U/ml and 5 mg/ml respectively were used throughout the previous steps. Nuclei were finally resuspended in “nuclear PBS” (0.7x PBS, 2mM MgCl2) containing 0.5% BSA and 0.2 U/ml RNase inhibitor. DAPI-stained nuclei were counted using a hemocytometer, and then were subject to fixation using the Parse Bioscience Evercode Fixation Kit (SB1003). Nuclei in DMSO were slow-frozen in a -80oC overnight according to the kit manual. A bar-coded snRNA-seq library was prepared using the Parse WT Mini Kit (ECW01010) and sequenced at the University of California, Irvine Genomics Research and Technology Hub.

#### Chromatin Immunoprecipitation Analysis

##### ChIP-qPCR

Percent input was calculated according to (Lin et al., 2012), where Input % = 100/2^([Cp[ChIP] – (Cp[Input] – Log2(Dilution Factor)). **ChIP-seq peak calling and IDR**: Reads were aligned to the *X. tropicalis* genome V10.0 using Bowtie 2 v2.4.1 (Langmead and Salzberg, 2012) using default options. The .sam alignment was converted to .bam, duplicates were removed and files were converted to .bed format using samtools (Li et al., 2009). Peaks were called against stage specific input DNA (Charney et al., 2017) using MACS2 v2.2.7.1 (Zhang et al., 2008) with the “-p .001” argument as the only non-default option. Irreproducibility discovery rate (IDR) was followed according to (Li et al., 2011), where optimal peaks were selected using p-value thresholds of 0.05 for true biological replicates, and 0.01 for pseudoreplicates of each individual biological replicate. **Heatmap generation**: DeepTools (Ramirez et al., 2014) “computeMatrix” was used to generate a matrix by mapping .bigwig coverage files to .bed peak regions, then “plotHeatmap” was used to visualize the matrix. Average signal intensity profiles (Fig2E, Supp1B, Fig5A) were made from matrices using “plotProfile”. **Motif analysis:** HOMER (V4.11) (Heinz et al., 2010) was used to predict de novo motifs using the “findMotifsGenome.pl” command within 100bp of the peak summit. To analyze motif occurrences within Foxi2 peaks, the following command was used: “annotatePeaks.pl $PEAKS $GENOME -size 2000 -hist 20 - ghist -m $MOTIF -mbed $MOTIF_BED > $OUTPUT”. The resulting matrix was visualized in Python using numpy (v1.26.2 (Harris et al. 2020)) and matplotlib (v3.8.2 (Hunter 2007)). **Peak location annotation**: To categorize whether a peak fell within a particular genomic region “annotatePeaks.pl $PEAKS $GENOME_FASTA -gtf $GFF3 > $OUTPUT” was used. **Genome browser visualization**: After alignment, .bam files were converted to .bed files used samtools command ‘bamtobed’. HOMER’s “makeTagDirectory” was then used to generate a tag directory as an input for “makeUCSCfile” which outputs a bedgraph. The Integrative Genome Viewer (IGV) was then used to convert the bedgraph into a .tdf file using igvtools “toTDF”. **Enhancer ranking**: The rank ordered super enhancer (ROSE) from the Young lab (Whyte et al., 2013) was used to categorically assign epigenetic peaks as regular enhancers (REs) or super enhancers (SEs) based on an enhancers over-all length and signal density. SEs were generated by stitching adjacent enhancer loci when loci passed a signal threshold. **Published ChIP-seq datasets:** Ep300 Stage 9, 10.5 (Hontelez et al., 2015), H3K4me1 Stage 8 (Gentsch et al., 2019), Stage 9, 10.5 (Hontelez et al., 2015), H3K27ac Stage 8, 9, 10.5 (Gupta et al., 2014). Animal and vegetal H3K4me1 at stage 10.25 (Paraiso et al., 2025).

#### Quantification and Statistical Analysis

##### RT-qPCR

The ΔΔCp method (Livak and Schmittgen, 2001) was utilized for calculating the fold-change in gene expression between treatment and control where the error among biological replicates was calculated using standard deviation and significance using a 2-tailed t-test. Bulk RNA-seq: Reads were aligned using RSEM V1.3.3 (Li and Dewey, 2011) with STAR V2.7.3 (Dobin et al., 2013) to the *Xenopus tropicalis* V10.0 genome to generate normalized expected read counts. Differential gene expression analysis was performed in R V4.1.1 using DEseq2 V1.34.0 (Love et al., 2014). **Localized gene expression analysis:** To determine ectoderm and endoderm localized genes, pre-existing data early gastrula tissue dissection RNA-seq data from (Blitz et al., 2017) was used. When comparing vegetal mass and animal cap dissection, genes with 2-fold enrichment in either germ layer were categorized as local, then the top 250 zygotic genes ranked by false discovery rate (FDR) were extracted **Zygotic gene expression timecourse:** To generate expression timecourses for zygotic gene sets, data from (Owens et al. 2016) was used. The 0HPF timepoint was used for detecting maternal transcript expression to segregate zygotically expressed genes in downstream timepoints. 0HPF timepoint TPMs were compared with downstream timepoint TPMs to calculate zygotic gene expression log fold-changes.

#### snRNA-seq Analysis

Barcode deconvolution was performed by Parse Biosciences. **Pre-processing:** Using scanpy v1.9.6 (Wolf et al. 2018), pandas v2.1.4 (McKinney et al. 2010), the nuclear data was log transformed and normalized and highly variable genes were annotated. Doublets were predicted using Scrublet (Wolock et al. 2019), then manually inspected, confirmed and removed based on illogical ectopic co-expression of germ-layer specific markers and outlier read counts. **Cell lineage annotation:** Leiden (Traag et al. 2019) was used to predict de novo cluster formation, visualized via Uniform Manifold Approximation Projection (UMAP) (McInnes et al. 2018). Scanpy was used to calculate differential gene expression analysis of known marker genes between de novo clusters which were assigned to their germ-layer identity. Accordingly, clusters were merged based on co-expression of known marker genes. In this way, cluster boundaries are initially formed in an unbiased way through de novo clustering, based on differential gene expression. *In-situ* hybridization studies of marker genes were subsequently used for appropriate cluster annotation and merger. **Gene expression analysis:** Pandas was used for both super enhancer genes’, and Foxi2/Sox3 associated genes’ z-score localization calculations by examining gene expression within the cells from each cell-type annotation, or merged annotations. Coefficient of Variation (COV) and average expression were calculated using numpy v1.26.2 (Harris et al. 2020).

## Acknowledgements

This work was made possible, in part, through access to the University of California, Irvine Genomics Research and Technology Hub (GRT Hub) parts of which are supported by NIH grants to the Comprehensive Cancer Center (P30CA-062203) and the UCI Skin Biology Resource Based Center (P30AR075047), as well as to the GRT Hub for instrumentation (1S10OD010794 and 1S10OD021718). We thank Xenbase for genomic and community resources (http://www.xenbase.org/, RRID: SCR_003280), and the University of California, Irvine High Performance Computing Cluster (https://hpc.oit.uci.edu/) for their valuable resources and helpful staff. This research was funded by the following grants awarded to K.W.Y.C. National Institute of Health R01 GM126395, R35 GM139617 and National Science Foundation 1755214. CLH is a recipient of a US Department of Education GAANN fellowship (P200A220015).

## Author contributions

C.L.H., I.L.B. and K.W.Y.C. designed the study, wrote the manuscript and performed experiments. A.H., J.C., and K.P. performed Ep300, H3K27ac, and H3K4me1 ChIP-seq; M.W.K. generated the Sox3 antibody and M.J.K. generated the Foxi2 antibody. C.L.H performed bioinformatics analyses.

## Declaration of interests

The authors declare no competing interests.

## Notes

### Competing Interest Statement

The authors have declared no competing interest.

