## Supplemental Figures and Legends for "Foxi2 and Sox3 are master regulators controlling ectoderm germ layer specification"

### Supp. Fig. 1

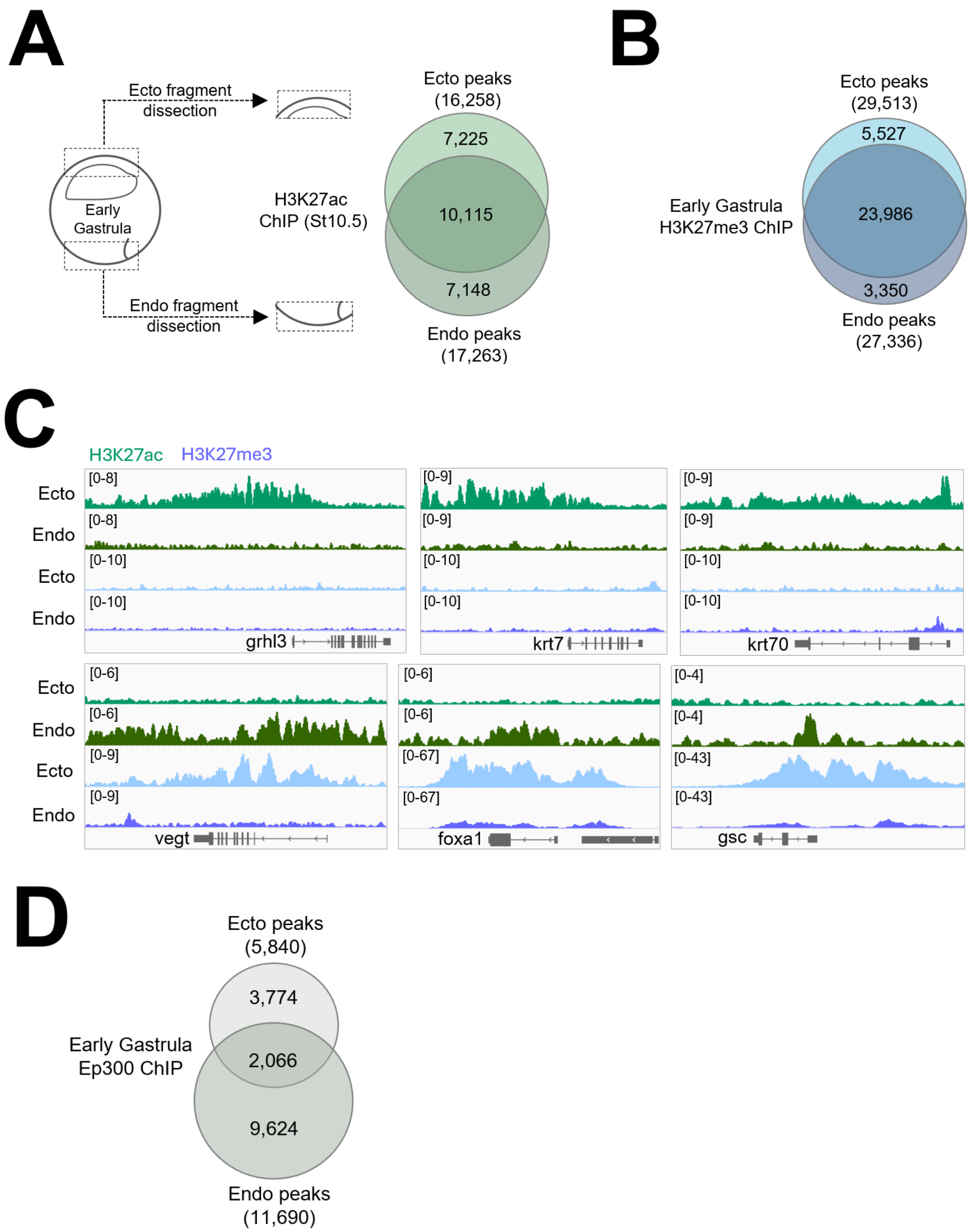
A
Ecto fragment dissection
Early Gastrula
Endo fragment dissection
H3K27ac ChIP (St10.5)
Ecto peaks (16,258)
7,225
10,115
7,148
Endo peaks (17,263)
B
Early Gastrula H3K27me3 ChIP
Ecto peaks (29,513)
5,527
23,986
3,350
Endo peaks (27,336)
C
H3K27ac H3K27me3
Ecto
Endo
Ecto
Endo
grhl3
krt7
krt70
vegt
foxa1
gsc
D
Early Gastrula Ep300 ChIP
Ecto peaks (5,840)
3,774
2,066
9,624
Endo peaks (11,690)

Supp. Fig. 2

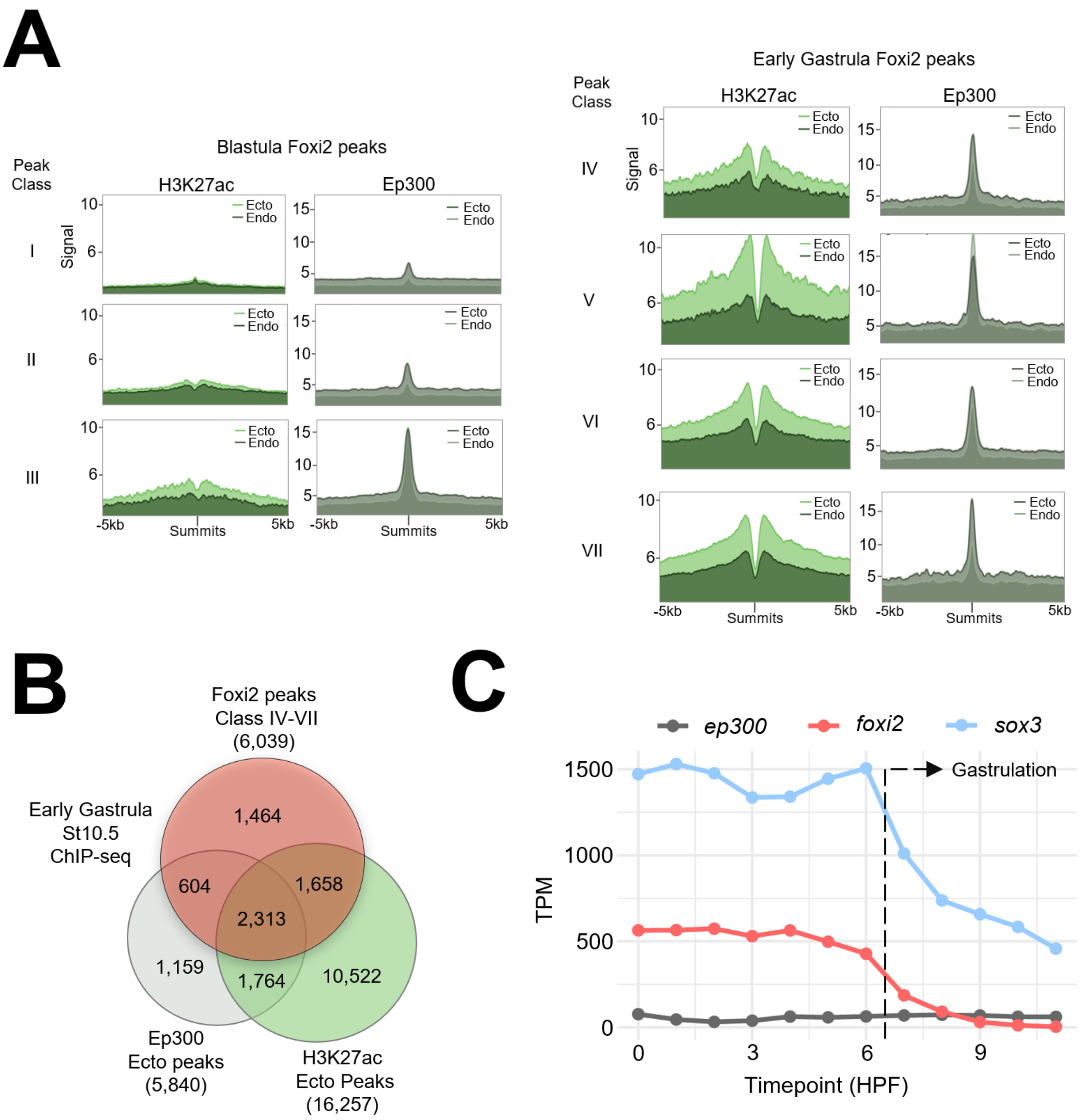

A
Blastula Foxi2 peaks
Peak Class
H3K27ac
Ep300
Signal
Ecto
Endo
I
II
III
-5kb
Summits
5kb
Early Gastrula Foxi2 peaks
IV
V
VI
VII
B
Foxi2 peaks
Class IV-VII
(6,039)
Early Gastrula
St10.5
ChIP-seq
1,464
604
1,658
2,313
1,159
1,764
10,522
Ep300
Ecto peaks
(5,840)
H3K27ac
Ecto Peaks
(16,257)
C
ep300
foxi2
sox3
Gastrulation
TPM
Timepoint (HPF)

### Supp. Fig. 3

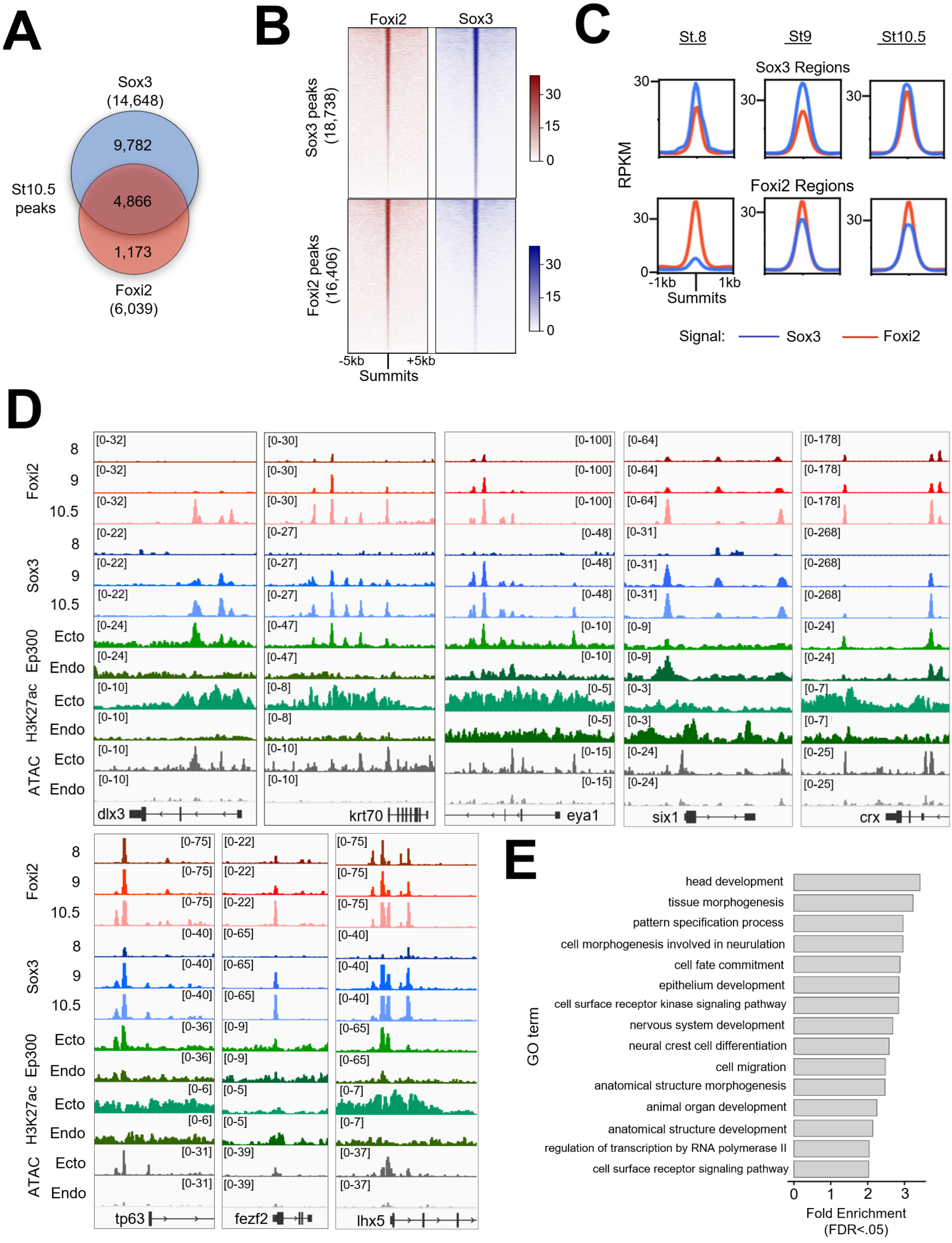

A
Sox3
(14,648)
9,782
4,866
1,173
St10.5
peaks
Foxi2
(6,039)
B
Foxi2
Sox3
Sox3 peaks
(18,738)
Foxi2 peaks
(16,406)
30
15
0
-5kb
+5kb
Summits
C
St.8
St9
St10.5
Sox3 Regions
Foxi2 Regions
RPKM
30
0
-1kb
1kb
Summits
Signal:
Sox3
Foxi2
D
Foxi2
Sox3
Ep300
H3K27ac
ATAC
8
9
10.5
Ecto
Endo
dlx3
krt70
eya1
six1
crx
tp63
fezf2
lhx5
E
GO term
head development
tissue morphogenesis
pattern specification process
cell morphogenesis involved in neurulation
cell fate commitment
epithelium development
cell surface receptor kinase signaling pathway
nervous system development
neural crest cell differentiation
cell migration
anatomical structure morphogenesis
animal organ development
anatomical structure development
regulation of transcription by RNA polymerase II
cell surface receptor signaling pathway
0
1
2
3
Fold Enrichment
(FDR<.05)

### Supp. Fig. 4

**A**

Foxi2 MO & Rescue Construct

5’CTGTGAATGTCCCACCAGACTTCATAAAGGATTATTATGAACACTTT WT *foxi2*

3’CACTTACAGGGTGGTCTGAAGTATT5’ *foxi2* MO

5’CGAATTCCCATGGACTACAAGGACGACGACGACAAGGGGAACACTTT *foxi2* Rescue

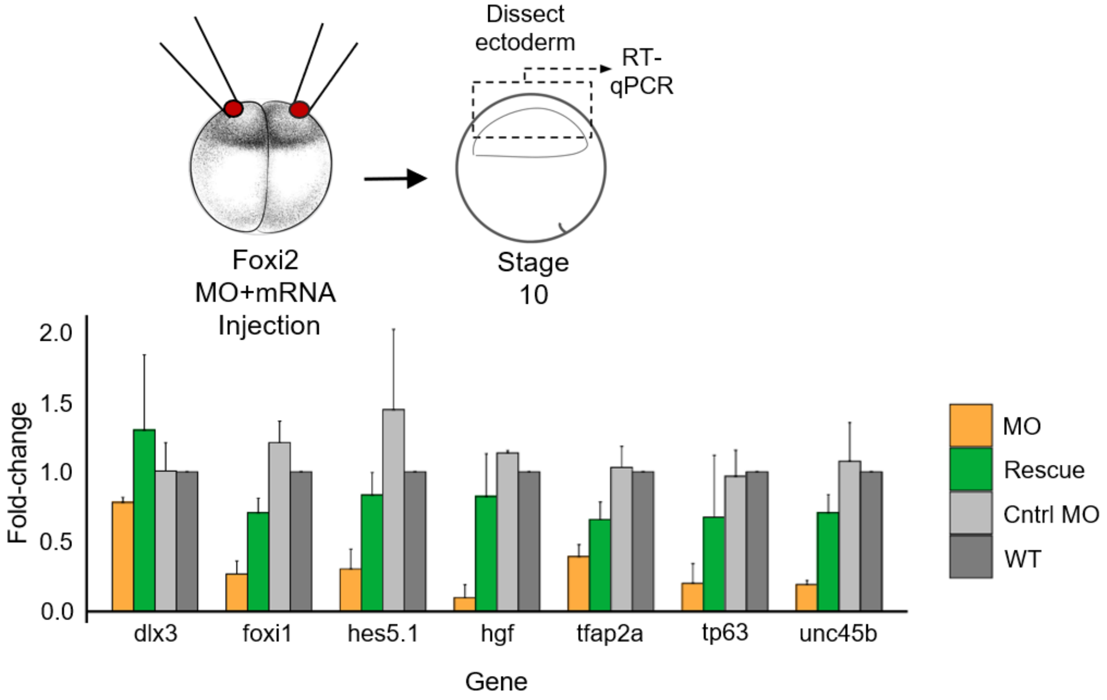


**B**

Sox3 MO & Rescue Construct

5’CCAGATGTATAGCATGTTGGACACAGACCTCAAGAGC3’ WT *sox3*

3’CTACATATCGTACAACCTGTGTCT 5’ *sox3* MO

5’TTTGGATCCGCCACCATGTACAGTATGCTGGATACTGATCTCAAGAGCCCGGTG *sox3* Rescue

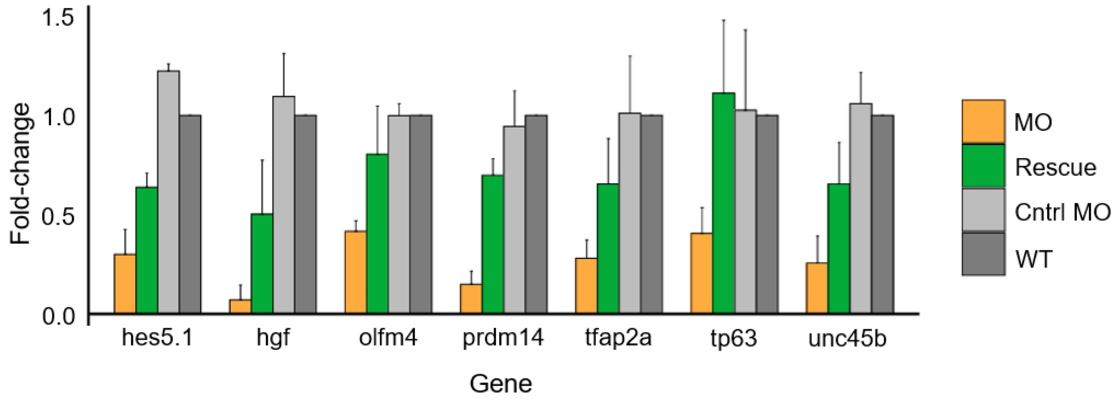

### Supp. Fig. 5

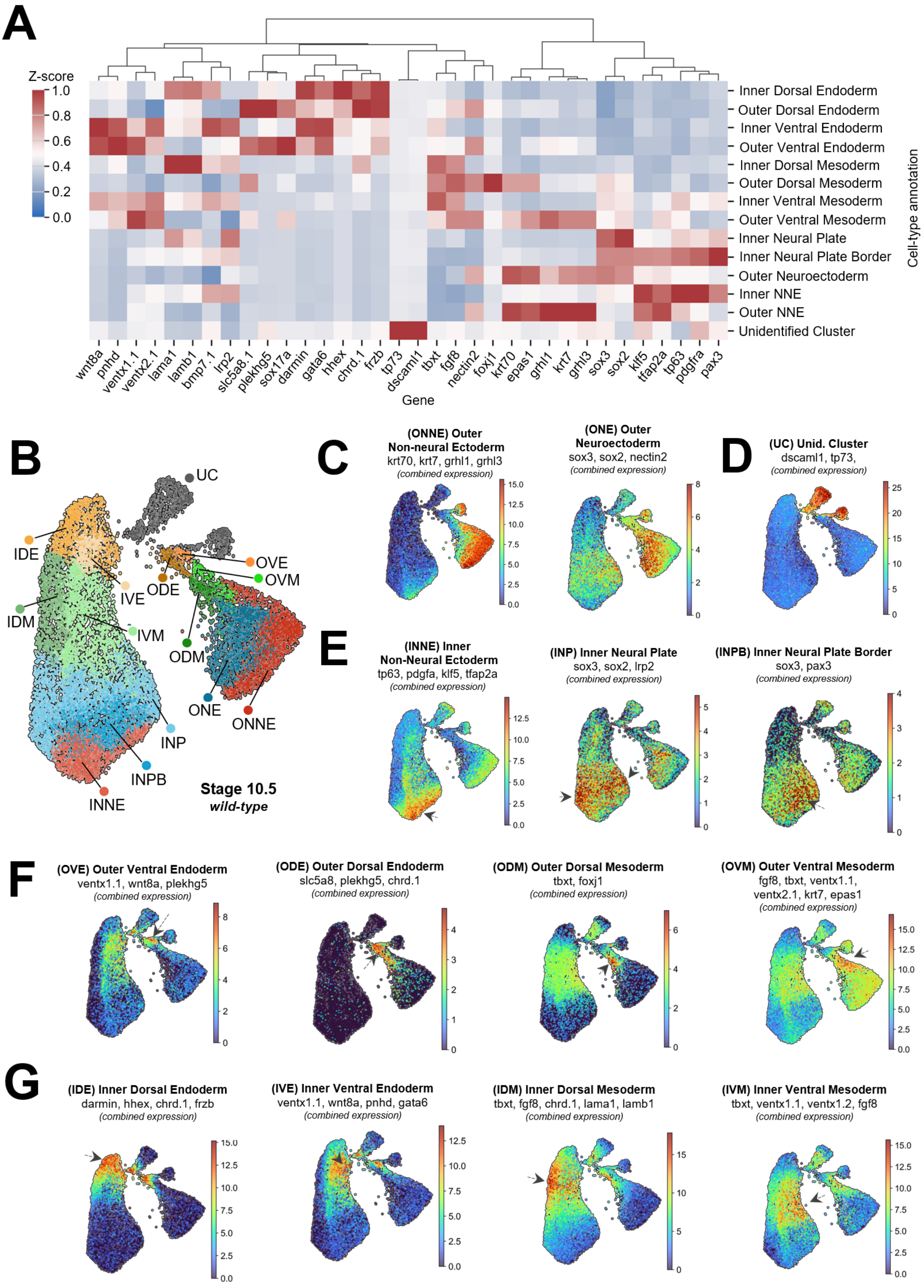

A
Z-score
1.0
0.8
0.6
0.4
0.2
0.0
Inner Dorsal Endoderm
Outer Dorsal Endoderm
Inner Ventral Endoderm
Outer Ventral Endoderm
Inner Dorsal Mesoderm
Outer Dorsal Mesoderm
Inner Ventral Mesoderm
Outer Ventral Mesoderm
Inner Neural Plate
Inner Neural Plate Border
Outer Neuroectoderm
Inner NNE
Outer NNE
Unidentified Cluster
Cell-type annotation
wnt8a pnhd ventx1.1 ventx2.1 lama1 lamb1 bmp7.1 lrp2 slc5a8.1 plekhg5 sox17a darmin gata6 hhex chrd.1 frzb tp73 dscaml1 tbxt fgf8 nectin2 foxj1 krt70 epas1 grhl1 krt7 grhl3 sox3 sox2 klf5 tfap2a tp63 pdgfra pax3
Gene
B
UC
IDE
IVE
ODE
OVE
OVM
IDM
IVM
ODM
ONE
ONNE
INP
INPB
INNE
Stage 10.5
wild-type
C
(ONNE) Outer Non-neural Ectoderm
krt70, krt7, grhl1, grhl3
(combined expression)
(ONE) Outer Neuroectoderm
sox3, sox2, nectin2
(combined expression)
D
(UC) Unid. Cluster
dscaml1, tp73,
(combined expression)
E
(INNE) Inner Non-Neural Ectoderm
tp63, pdgfa, klf5, tfap2a
(combined expression)
(INP) Inner Neural Plate
sox3, sox2, lrp2
(combined expression)
(INPB) Inner Neural Plate Border
sox3, pax3
(combined expression)
F
(OVE) Outer Ventral Endoderm
ventx1.1, wnt8a, plekhg5
(combined expression)
(ODE) Outer Dorsal Endoderm
slc5a8, plekhg5, chrd.1
(combined expression)
(ODM) Outer Dorsal Mesoderm
tbxt, foxj1
(combined expression)
(OVM) Outer Ventral Mesoderm
fgf8, tbxt, ventx1.1, ventx2.1, krt7, epas1
(combined expression)
G
(IDE) Inner Dorsal Endoderm
darmin, hhex, chrd.1, frzb
(combined expression)
(IVE) Inner Ventral Endoderm
ventx1.1, wnt8a, pnhd, gata6
(combined expression)
(IDM) Inner Dorsal Mesoderm
tbxt, fgf8, chrd.1, lama1, lamb1
(combined expression)
(IVM) Inner Ventral Mesoderm
tbxt, ventx1.1, ventx1.2, fgf8
(combined expression)

### Supp. Fig. 6

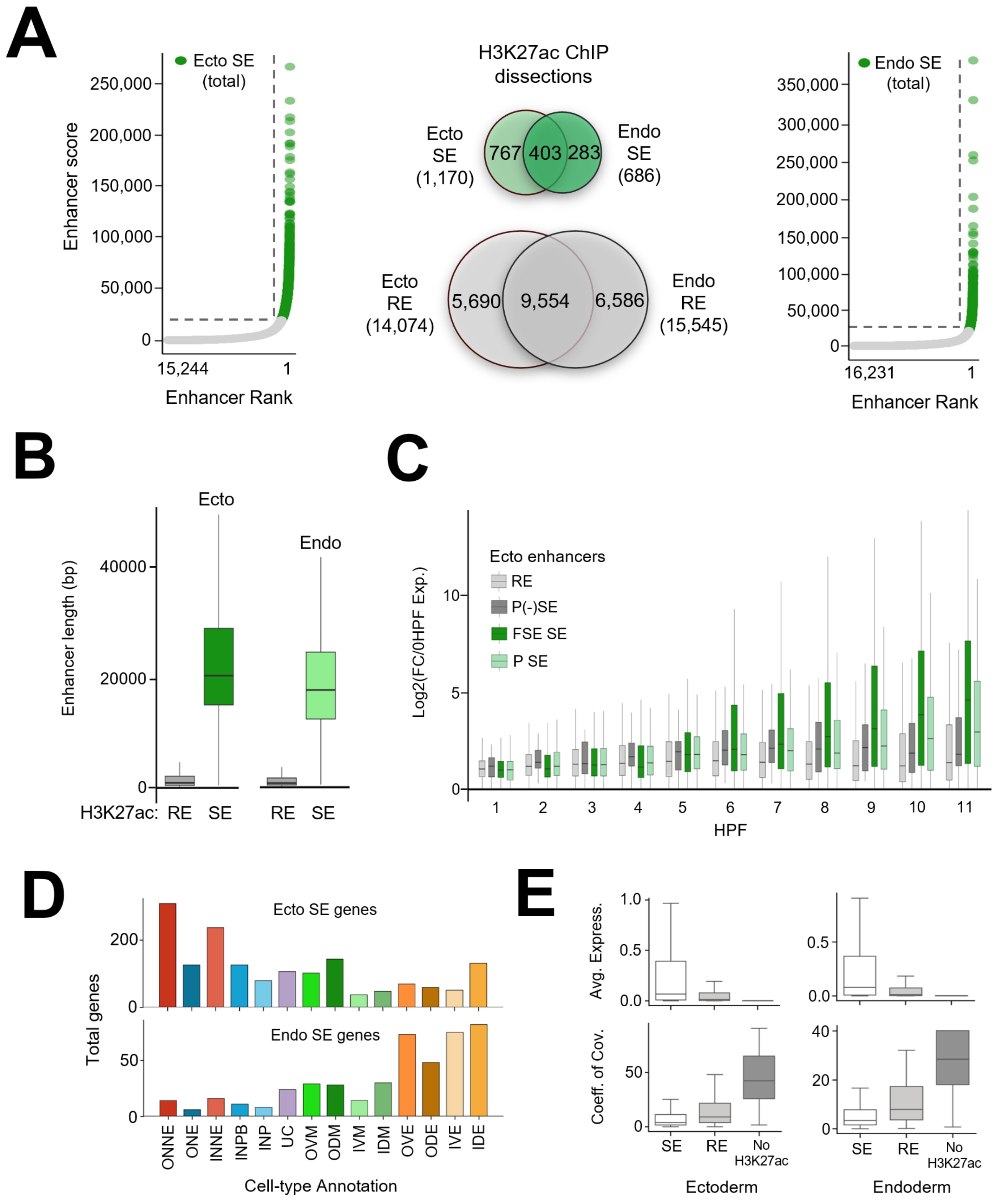
A
Ecto SE (total)
Enhancer score
250,000
200,000
150,000
100,000
50,000
0
15,244
1
Enhancer Rank
H3K27ac ChIP dissections
Ecto SE (1,170)
767
403
283
Endo SE (686)
Ecto RE (14,074)
5,690
9,554
6,586
Endo RE (15,545)
Endo SE (total)
350,000
300,000
250,000
200,000
150,000
100,000
50,000
0
16,231
1
Enhancer Rank
B
Ecto
Endo
Enhancer length (bp)
40000
20000
0
H3K27ac: RE SE RE SE
C
Ecto enhancers
RE
P(-)SE
FSE SE
P SE
Log2(FC/0HPF Exp.)
10
5
0
1 2 3 4 5 6 7 8 9 10 11
HPF
D
Ecto SE genes
Endo SE genes
Total genes
200
0
50
0
ONNE ONE INNE INPB INP UC OVM ODM IVM IDM OVE ODE IVE IDE
Cell-type Annotation
E
Avg. Express.
1.0
0.5
0.0
Coeff. of Cov.
50
0
40
20
0
SE RE No H3K27ac
Ectoderm
Endoderm

#### Supplementary Figure Legends

Figure S1. *Epigenetic Landscape of Early Gastrula Ectoderm and Endoderm.* (A) Schematic depicting early gastrula ectoderm and endoderm dissections performed for ChIP-seq (left). Total H3K27ac peaks detected in ectoderm and endoderm dissections (right). (B) Total H3K27me3 peaks detected in ectoderm and endoderm dissections. (C) Genome browser views showing differential epigenetic signatures of key ectoderm and endoderm expressed genes. (D) Total Ep300 peaks detected in ectoderm and endoderm dissections.

Figure S2. *Spatial and Temporal Epigenetic Regulation of Foxi2 Peaks in Early Gastrula.* (A) H3K27ac and Ep300 signal in ectoderm and endoderm across Foxi2 peak classes I-III (blastula stage, left) and VI-VII (early gastrula stage, right). (B) Intersection of early gastrula Foxi2 peaks and ectoderm dissected H3K27ac and Ep300 peak regions. (C) Temporal expression profiles of maternal factors during early embryogenesis.

Figure S3. *Spatial and Temporal Binding Profiles of Foxi2 and Sox3*. (A) Early gastrula Foxi2 and Sox3 peak overlap. (B) Total binding profiles of Foxi2 and Sox3 peak. (C) Stage specific binding profiles of Foxi2 and Sox3 in respective peak regions. (D) Genome browser views of key neural and epidermal ectodermal genes. (E) GO annotation of genes near shared Foxi2, Sox3 and Ep300 peaks.

Figure S4. *Foxi2 and Sox3 Morphant Rescue: Design and Validation.* (A) Schematic depicting Foxi2 morpholino design (top) and RT-qPCR of Foxi2 morphant rescue in ectoderm dissections (bottom). (B) Schematic depicting Sox3 morpholino design (top) and RT-qPCR of Sox3 morphant rescue of ectoderm targets (bottom).

Figure S5. *Cell-Type Specific Marker Expression and UMAP Analysis of Early Gastrula Single-Nucleus RNA-Seq Data* (A) z-score analysis of early gastrula gene marker expression across annotated cell-types. (B) UMAP visualization of early gastrula snRNA-seq data with annotated cell types. (C) Combined expression UMAPs of outer ectoderm cell types and their markers (ONNE, ONE). (D) Combined expression UMAP of an unidentified cluster (UC). (E) Combined expression UMAPs of inner ectoderm cell types and their markers (INNE, INP, INPB). (F) combined expression UMAPs of outer mesendoderm cell types and their markers (OVE, ODE,

ODM, OVM). (G) Combined expression UMAPs of inner mesendoderm cell types and their markers (IDE, IVE, IDM, IVM).

Figure S6. *Comparative Analysis of Regular and Super Enhancers in Ectoderm and Endoderm:* (A) Rank Ordered Super-Enhancer (ROSE) calls: H3K27ac regular enhancer (RE) and super enhancer (SE) region overlaps between ectoderm (left) and endoderm (right) dissections. (B) Comparison of H3K27ac RE and SE lengths in ectoderm (left) and endoderm (right). (C) Temporal expression profiles of genes located within 20kb of ectoderm REs and SEs. (D) Cell type-specific z-score localization of genes located within 20kb of ectoderm or endodermal SEs. (E) Expression profiles of ectodermal and endodermal enhancer associated genes (top) and their coefficient of variation in expression (bottom).
